# Exact decoding of the sequentially Markov coalescent

**DOI:** 10.1101/2020.09.21.307355

**Authors:** Caleb Ki, Jonathan Terhorst

## Abstract

In statistical genetics, the sequentially Markov coalescent (SMC) is an important framework for approximating the distribution of genetic variation data under complex evolutionary models. Methods based on SMC are widely used in genetics and evolutionary biology, with significant applications to genotype phasing and imputation, recombination rate estimation, and inferring population history. SMC allows for likelihood-based inference using hidden Markov models (HMMs), where the latent variable represents a genealogy. Because genealogies are continuous, while HMMs are discrete, SMC requires discretizing the space of trees in a way that is complicated and can lead to bias. In this work, we propose a method that circumvents this requirement, enabling SMC-based inference to be performed in the natural setting of a continuous state space. We derive fast, exact methods for frequentist and Bayesian inference using SMC. Compared to existing methods, ours requires minimal user intervention or parameter tuning, no numerical optimization or E-M, and is faster and more accurate.

## 1. Introduction

Probabilistic models of evolution have played a central role in genetics since the inception of the field a century ago. Beginning with foundational work by Fisher (1930) and Wright (1931), and continuing with important contributions from Moran (1958), Kimura (1955a,b), Kingman (1982a,b,c), Griffiths (1981), and Hudson (1983), and many others, a succession of increasingly sophisticated stochastic models were developed to describe patterns of ancestry and genetic variation found in a population. Statisticians harnessed these models to analyze genetic data, initially with the now quaint-seeming goal of understanding the evolution of a single gene. More recently, as next-generation sequencing has enabled the collection of genome-wide data from millions of people, interest has risen in methods for studying evolution using large numbers of whole genomes.

In this article, we study a popular subset of those methods which are likelihood-based; that is, these methods work by inverting a statistical model that maps evolutionary parameters to a probability distribution over genetic variation data. As we will see, exact inference in this setting is impossible owing to the need to integrate out a high-dimensional latent variable which encodes the genome-wide ancestry of every sampled individual. Consequently, a number of approximate methods have been proposed, which try to strike a balance between biological realism and computational tractability.

We focus on one such approximation known as the *sequentially Markov coalescent* (SMC; McVean and Cardin, 2005; Marjoram and Wall, 2006; Carmi et al., 2014; Hobolth and Jensen, 2014). SMC^1^ assumes that the sequence of (random) genealogies at each position in the genome forms a Markov chain, thereby enabling efficient likelihood-based inference using hidden Markov models (HMMs). Although the Markov assumption is wrong (Wiuf and Hein, 1999), it has nevertheless proved highly useful in practice. In particular, both the influential haplotype copying model of Li and Stephens (2003) and the popular program PSMC (Li and Durbin, 2011) for inferring population history are SMC methods.

In order to bring the HMM machinery to bear on this problem, additional and somewhat awkward assumptions are needed. The latent variable in an HMM must have finite support, whereas the latent variable in SMC is a continuous tree. Therefore, the space of trees must be discretized, and, in some cases, restrictions must also be placed on the topology of each tree. In applications, the user must select a discretization scheme, a non-obvious choice which nonetheless has profound consequences for downstream inference (Parag and Pybus, 2019). The main message of our paper is that this is not necessary: it is possible to solve the SMC exactly, in its natural setting of continuous state space. We accomplish this by slightly modifying the standard SMC model in a way that does not impact inference, but renders the problem theoretically and computationally much easier. In particular, this modification allows us to leverage recent innovations in changepoint detection, leading to algorithms which not only have less bias than existing approaches, but also outperforms them computationally.

The rest of the paper is organized as follows. In Section 2 we formally define our data and model, introduce notation, and survey related work. In Section 3 we derive our main results: exact and efficient Bayesian and frequentist algorithms for inferring genealogies from genetic variation data. In Section 4 we thoroughly benchmark our method, compare it to existing approaches, and provide an application to real data analysis. We provide concluding remarks in Section 5.

## 2. Background

In this section we introduce notation, formalize the problem we want to solve, and survey earlier work. We presume some familiarity with standard terminology and models in genetics; introductory texts include Hein, Schierup and Wiuf (2005) and Durrett (2008).

### 2.1. Motivation

Our method aims to infer a sequence of latent genealogies using genetic variation data. To motivate our interest in this, consider first a related problem with a more direct scientific application: given a matrix of DNA sequence data **Y** ∈ {`A`, `C`, `G`, `T}`^*H×N*^ from *H >* 1 homologous chromosomes each *N* base pairs long, and an evolutionary model *ϕ* hypothesized to have generated these data, find the likelihood *p*(**Y**|*ϕ*). This generic formulation encompasses a wide variety of inference problems in genetics and evolutionary biology; if we could easily solve it, important new scientific insights would result.

Unfortunately, this is not possible using current methods. The difficulty lies in the fact that the relationship between the data **Y** and the scientifically interesting quantity *ϕ* is mediated through a complex, latent combinatorial structure known as the ancestral recombination graph (ARG; Griffiths and Marjoram, 1997), which encodes the genealogical relationships between every sample at every position in the genome. The ARG is sufficient for *ϕ*: evolution generates the ARG, and conditional on it, the data contain no further information about *ϕ*. Thus, the likelihood problem requires the integration

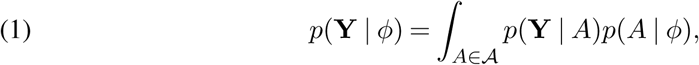

where *A* denotes an ARG, and 𝒜 denotes the support set of ARGs for a sample of *H* chromosomes. This is a very challenging integral; although a method for evaluating it is known (Griffiths and Marjoram, 1996), it only works for small data sets. That is because, for large *N* and *H*, there are a huge number of ARGs that could have plausibly generated a given data set, such that the complexity of 𝒜 explodes as *N* and *H* grow. Indeed, (1) cannot be computed for chromosome-scale data even for the simplest case *H* = 2.

The sequentially Markov coalescent addresses this problem by decomposing the ARG into a sequence of marginal gene trees *X*_1_, …, *X*_*N*_, one for each position in the chromosome, and supposing that this sequence is Markov. Then, we have

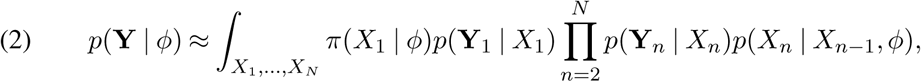

where *π*(·|*ϕ*) is a stationary distribution for the Markov chain *X*_1_, …, *X*_*N*_, *p*(*X*_*n*_|*X*_*n*−1_, *ϕ*) is a transition density, and [**Y**_1_|…| **Y**_*N*_] = **Y** are the data at each site. If the *X*_*i*_ have discrete support, then this represents a hidden Markov model, whence (2) can be efficiently evaluated using the forward algorithm. For estimating *ϕ*, E-M type algorithms are generally preferred, and these require computing the posterior distribution *p*(*X*_1_, …, *X*_*N*_ | **Y**, *ϕ*).

### 2.2. Demographic inference

To make this problem more concrete, in this paper we focus specifically on computing (1) when the chromosomes evolve under selective neutrality, and *ϕ* represents historical fluctuations in population size. In this case, we can identify *ϕ* with a function *N*_*e*_ : [0, ∞) → (0, ∞), such that *N*_*e*_(*t*) was the coalescent effective population size *t* generations before the present (Durrett, 2008, §4.4). This function governs the marginal distribution of coalescence time at a particular locus in a sample of two chromosomes. Specifically, setting *η*(*t*) = 1*/N*_*e*_(*t*), the density of this time is

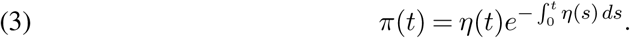

Note that *η*(*t*) = 1 recovers the well-known case of Kingman’s coalescent, *π*(*t*) = *e*^−*t*^, which we treat as the default prior in what follows.

Apart from intrinsic interest in learning population history, it is important to get a sharp estimate of *N*_*e*_(*t*) as unmodeled variability in *N*_*e*_(*t*) confound attempts to study other evolutionary phenomena such as natural selection, or mutation rate variation. Estimation of this function is known in the literature as *demographic inference* (Spence et al., 2018). For the remainder of the paper we will focus on this application. To simplify the notation, we suppress explicit dependence on *N*_*e*_(*t*) and capture it implicitly through the function *π*, and we even suppress dependence on *π* when it is clear from context.

### 2.3. Our contribution

As discussed in Section 1, discretizing *X*_*i*_ is unnatural and results in bias. In this work, we derive efficient methods for computing the posterior distribution *p*(*X*_1_, …, *X*_*N*_ | **Y**), or its *maximum a posteriori* estimate

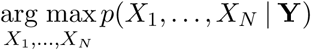

for a given demography *π*, when each *X*_*i*_ is a tree with continuous branch lengths. That is, unlike existing methods, we do not assume that the set of possible *X*_*i*_ is discrete or finite. For the important case of *H* = 2 chromosomes, our method is “exact” in the sense that it is devoid of further approximations (beyond the standard ones which we outline in the next section). For *H >* 2 our method makes additional assumptions about the topology of each *X*_*i*_, but still retains the desirable property of operating in continuous time.

### 2.4. Notation and model

We now fix necessary notation and define the model that is used to prove our results. For ease of exposition, our results focus on the simplest possible case of analyzing a pair of chromosomes (*H* = 2 in the notation of the previous section). In Section 3.4 we describe how to extend our results to larger sample sizes

Assume that that we have sampled a pair of homologous chromosomes each consisting of *N* non-recombining loci. Meiotic recombination occurs between loci with rate *ρ* per unit time, and does not occur within each locus.^2^ The number generations backwards in time until the two chromosomes meet at a common ancestor (TMRCA) at locus *i* is denoted *X*_*i*_ ∈ ℝ_*>*0_. The number of positions where the two chromosomes differ at locus *i* is denoted by *Y*_*i*_. Under a standard assumption known as the infinite sites model (Durrett, 2008, §1.4), *Y*_*i*_ has the conditional distribution

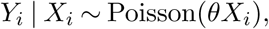

where *θ* is the mutation rate. We assume that both *θ* and *ρ* are small. In particular, some of our proofs rely on the fact that *ρ* ≪ 1. These are fairly mild assumptions which hold in many settings of interest. For example, in humans, the population-scaled rates of mutation and recombination per nucleotide are around 10^−4^. Conversely, if recombinations are frequent, then there is little advantage in employing the methods we describe here, which depend on the presence of linkage disequilibrium between nearby loci.

The sequentially Markov coalescent is a generative model for the sequence *X*_1_, …, *X*_*N*_, which we abbreviate as *X*_1:*N*_ henceforth (and similarly for *Y*_1:*N*_). SMC characterizes how shared ancestry changes when moving from one locus to the next. Assuming there is at most one recombination between adjacent loci, and we can specify an SMC model by the conditional density

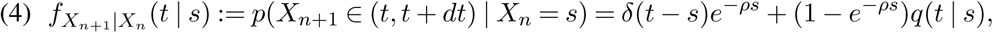

where *δ*() is the Dirac delta function, and *q*(*t*|*s*) is the conditional density of *t* given that a recombination occurred and that the existing TMRCA equals *s*. Various proposals for *q*(*t*|*s*) exist in the literature, each with slightly different properties (McVean and Cardin, 2005; Marjoram and Wall, 2006; Li and Durbin, 2011; Hobolth and Jensen, 2014; Carmi et al., 2014). Importantly, they share the common feature that (4) is (approximately, in the case of Li and Durbin, 2011) reversible with respect to the coalescent. That is,

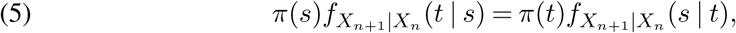

where the stationary measure *π* was defined in equation (3).

### 2.5. Connection to changepoint detection

Our work is motivated by the observation that (4) is *almost* a changepoint model. Indeed, SMC can be viewed as a prior over the space of piecewise constant functions spanning the interval [0, *N*); conditional on realizing one such function, say *ξ* : [0, *N*) → [0, ∞), each *X*_*i*_ = *ξ*(*i* − 1), and the data *Y*_1:*N*_ are independent Poisson draws with mean 𝔼(*Y*_*i*_ | *X*_*i*_) = *θX*_*i*_. In genetics, each contiguous segment where *X*_*i*_ = *X*_*i*+1_ = … = *X*_*i*+*k*−1_ = *τ*, say, is known as an *identity by descent* (IBD) tract, with *time to most recent common ancestor* (TMRCA) *τ*; the flanking positions where *X*_*i*−1_ ≠ *X*_*i*_ and *X*_*i*+*k*_ ≠ *X*_*i*+*k*−1_ are called *recombination breakpoints* (e.g., Browning and Browning, 2011). In changepoint detection, these are called *segments, segment heights* (or just heights), and *changepoints*, respectively. In what follows, we use these terms interchangeably depending on what is most descriptive in a given context.

A standard assumption in changepoint detection is that neighboring segment heights are independent, which is to say that *X*_*i*_ ⊥ *X*_*i*+1_ for any *i* such that *X*_*i*_ ≠ *X*_*i*+1_. As we will see, this enables fast and accurate algorithms for inferring the sequence *X*_1:*N*_. SMC violates this assumption through the conditional density *q*(*t*|*s*): the correlation between *t* and *s* in (4) makes the problem non-standard from a changepoint perspective. It is tempting to simply ignore it. Indeed, if *q*(*t*|*s*) were replaced by some function *<u>π</u>* (*t*) which did not depend on *s*, then (4) would become a so-called product partition model (PPM; Barry and Hartigan, 1992). PPMs are well-understood. In particular, efficient methods have been developed to analyze PPMs in both Bayesian (Barry and Hartigan, 1993; Fearnhead, 2006) and frequentist (Jackson et al., 2005; Killick, Fearnhead and Eckley, 2012) settings.

### 2.6. A renewal approximation

In biological applications, the orientation of the data sequence *Y*_1:*N*_ is arbitrary; we could equivalently work with the reversed sequence *Y*_*N*_, *Y*_*N*−1_, …, *Y*_1_ instead. Additionally, both theoretical and empirical evidence overwhelmingly support that Kingman’s coalescent is a robust and accurate description of ancestry at a particular gene. For these reasons, it is important that any SMC model maintain the detailed balance condition (5). Given this desideratum, the obvious choice for *<u>π</u>* becomes

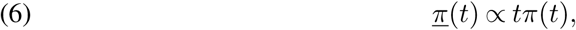

leading to the modified transition density

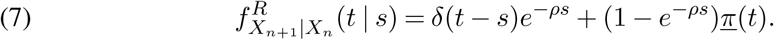

Checking the detailed balance condition (5), we obtain

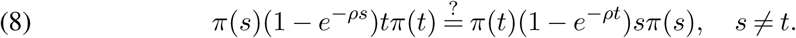

Though (8) is not true in general, equality holds when both sides are expanded to first-order in *ρ*, which suffices for the applications we consider here. So, we will assume henceforth that *ρ* is sufficiently small that (8) holds as an identity.

The renewal approximation preserves an important piece of prior information concerning the nature of identity-by-descent. Recall that *<u>π</u>* as defined in (6) is the so-called *length-biased* distribution corresponding to *π* (Feller, 1971). Length bias emerges precisely because of the level-dependent nature of IBD tract lengths under SMC: given an IBD tract with TMRCA *x*, the rate of recombination is *ρx*, so more recent IBD tracts are longer. Therefore, a randomly sampled location on the chromosome is more likely to fall on a longer tract and be recent. *<u>π</u>* “undoes” this bias and restores stationarity with respect to *π*.

Thus, compared with the standard SMC model in (4), the modified formulation in (7) retains prior information on the dependence between IBD segment length and height, while dropping prior information on the correlation between neighboring segment heights. We hypothesized that, for inference, it is more important that the prior capture the former effect than the latter. This is similar to the observation in changepoint detection that identifying changepoint locations tends to be harder than identifying the corresponding segment heights. Conditional on a given segmentation, finding the most likely segment heights is usually trivial, with a solution that depends mostly on the data and very little on the prior. Thus, it seems most important to encode prior information about the nature of the segmentation itself.

### 2.7. Prior work

The Markov chain defined by (7) was previously studied by Carmi et al. (2014), who coined the term renewal approximation. Carmi et al. derived theoretical results and performed simulations to study identity-by-descent patterns produced by SMC models. They found that the renewal approximation is comparable to other variants of SMC with some inaccuracy mainly in the tails of the IBD distribution. Importantly, these results pertain to the accuracy of these methods as *priors*; they do not necessarily imply that the renewal approximation is inferior for *inference*. Indeed, generally one hopes that “the data overwhelm the prior,” so that inferences do not depend strongly on the choice of prior model. We hypothesized that the ability to analyze significantly larger quantities of data, with less bias, would outweigh any penalty incurred through the use of a more approximative prior.

There have been a few papers specifically devoted to improving the efficiency of SMC. The naive forward-backward and Viterbi algorithms for HMMs take *O*(*LM* ^2^) time when applied to SMC, where *L* is the length of the analyzed sequence, and *M* is the number of hidden states (time discretizations) used to approximate the coalescent state space. By exploiting the specialized structure of the SMC transition matrix, Harris et al. (2014) were able to reduce this to *O*(*ML*) for the SMC model of McVean and Cardin (2005). Palamara et al. (2018) extended these results to the so-called SMC’ model of Marjoram and Wall (2006). In a different line of work, Lunter (2019) recently showed that MAP estimation can be performed for the Li and Stephens model in *O*(*L*) time irrespective of the size *H* of the underlying copying panel, after a preprocessing step that costs *O*(*HL*) time (Durbin, 2014). Compared to these works, we will show that our method has empirical running time *O*(*HL*), without requiring discretization or making strong genealogical assumptions as in the Li and Stephens model.

More generally, SMC is the foundation of a large number of other inference methods in genetics. Haplotype copying models (Li and Stephens, 2003) have been used to study natural selection (Voight et al., 2006), ancestry (Price et al., 2009), population structure (Lawson et al., 2012), and population history (Gay, Myers and McVean, 2007); and to perform haplotype phasing and imputation (Scheet and Stephens, 2006; Marchini et al., 2007; Howie, Donnelly and Marchini, 2009). Similarly, PSMC and related methods for inferring population size history (Li and Durbin, 2011; Schiffels and Durbin, 2014; Terhorst, Kamm and Song, 2017; Steinrücken et al., 2019) are now a standard component of population genetic analysis, and have been cited in thousands of papers.

## 3. Methods

In this section we derive exact representations for the sequence of marginal posterior distributions *p*(*X*_*n*_ | *Y*_1:*N*_), *n* = 1, 2, …, *N*, and efficient algorithms for sampling paths from the posterior density *p*(*X*_1:*N*_ | *Y*_1:*N*_) and for computing the MAP path

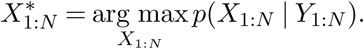

### 3.1. Exact marginal posterior

In what follows, we write *f* (*x*) ∈ ℳ_Γ_(*K*) to signify a the probability density *f* is a mixture of *K* gamma distributions, with the mixing weights, scale and shape parameters left unspecified. By abuse of notation, we also write *X* ∼ ℳ_Γ_(*K*) to signify that the random variable *X* is distributed according to such a mixture.

Let *α*(*X*_*n*_) = *p*(*X*_*n*_|*Y*_1:*n*_) denote the (rescaled) forward function from the standard forward-backward algorithm for inferring hidden Markov models (Bishop, 2006, §13.2.4). Our first result shows that, under the renewal approximation, *α*(*X*_*n*_) is a mixture of gamma distributions.

#### Lemma 1.

*Suppose that π*(*x*) ∈ ℳ_Γ_(*K*). *Then α*(*X*_*n*_) = *p*(*X*_*n*_ | *Y*_1:*n*_) ∈ ℳ_Γ_(*nK*).

The proof of Lemma 1 requires only a few simple facts from Bayesian analysis.

Fact 1. If *X* ∼ Γ(*a, b*) and *Y* | *X* ∼ Poisson(*θX*), then *X* | *Y* ∼ Γ(*a* + *Y, b* + *θ*).

Fact 2. If *X* ∼ ℳ_Γ_(*K*) and *Y* | *X* ∼ Poisson(*X*), then *X* | *Y* ∼ ℳ_Γ_(*K*).

Fact 3. If *X*_*n*_ | *Y*_*n*_ ∼ ℳ_Γ_(*nK*) and *π* ∈ ℳ_Γ_(*K*), then under the renewal approximation (7), *X*_*n*+1_ | *Y*_*n*_ ∼ ℳ_Γ_(*n* + 1)*K*).

The first two facts are well-known consequences of conjugacy. To establish the third, note that

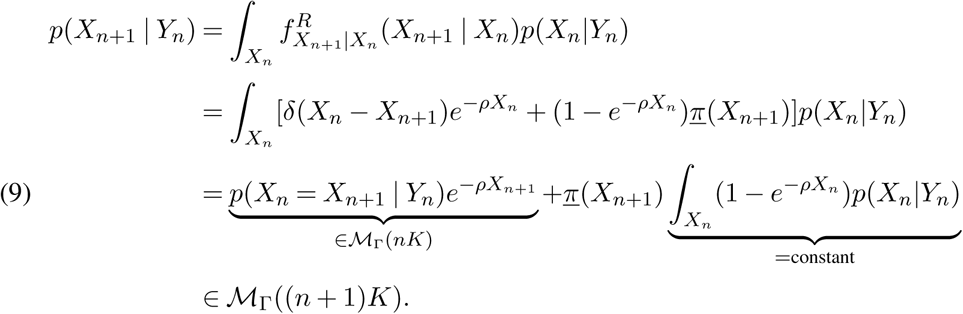

#### Proof of Lemma 1.

By induction on *n*. The case *n* = 1 follows from Facts 1 and 2. And, if the claim holds for *n* = *i*, then *X*_*i*+1_ ⊥ *Y*_1:*i*_ ∼ ℳ_Γ_((*i* + 1)*K*) by Fact 3. Since *Y*_*i*+1_ ⊥ *Y*_1:*i*_ | *X*_*i*+1_, Fact 2 implies

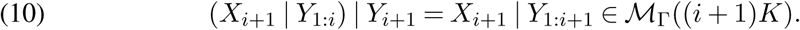

□

Using Lemma 1, we establish a representation theorem for *p*(*X*_*n*_ | *Y*_1:*N*_).

#### Theorem 1.

*If π*(*x*) ∈ *M*_Γ_(*K*) *then there exists f* (*X*_*n*_) ∈ ℳ_Γ_(*Kn*) *and g*(*X*_*n*_) ∈ ℳ_Γ_(*K*(*N* − *n*)) *such that*

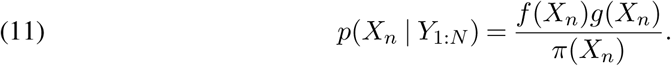

#### Proof.

Define 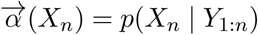 (*X*_*n*_) = *p*(*X*_*n*_ *Y*_1:*n*_) to be the quantity derived in Lemma 1, and let 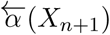 (*X*_*n*+1_) be obtained by running the forward algorithm from that lemma on the reversed sequence (*Y*_*N*_, *Y*_*N*−1_, …, *Y*_*n*+1_). By reversibility, 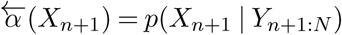 (*X*_*n*+1_) = *p*(*X*_*n*+1_ | *Y*_*n*+1:*N*_) and hence

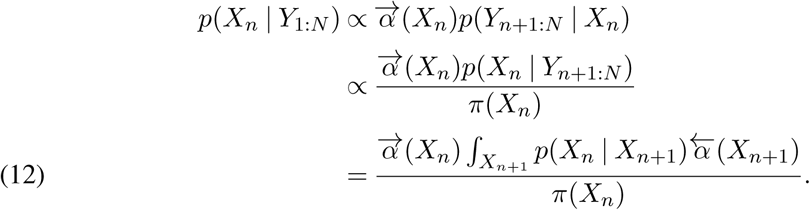

By Lemma 1, 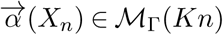 and 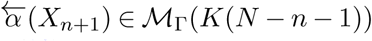. Finally, using the same argument that established equation (10),

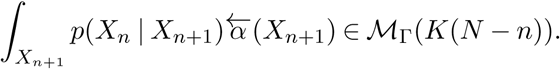

□

We can also derive exact expressions for the mixing proportions, shape, and scale parameters for *π*(*X*_*n*_)*p*(*X*_*n*_|*Y*_1:*N*_), as well as the correct normalizing constant. This requires substantial additional notation and is deferred to Appendix A.

#### Remark.

Instead of the reversibility argument used to prove Theorem 1, we could have used ideas from the proof of Lemma 1 to derive a sum-of-gammas representation for the rescaled backward function

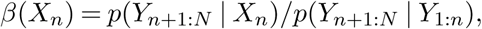

whence *p*(*X*_*n*_|*Y*_1:*N*_) = *α*(*X*_*n*_)*β*(*X*_*n*_). We experimented with this approach, but found that it was numerically unstable for long sequences: whereas the mixture coefficients of *α*(*X*_*n*_) live in the simplex, the backwards function *β*(*X*_*n*_) is not a probability distribution in *X*_*n*_, and we observed that the mixture coefficients tended to diverge when *N* was large. It seems that the rational representation (11) has superior numerical properties.

### 3.2. Efficient posterior sampling

The exact posterior formula derived in Theorem 1 is useful for visualization, or numerically evaluating functionals (e.g., the posterior mean) of the posterior distribution. However, it is less suited to sampling because:

1. The denominator does not divide the numerator except when *K* = 1, so the posterior is not a mixture in general; and
2. Even then, sampling requires expanding the numerator in (12) into (as many as) *O*(*K*^2^*N* ^2^) mixture components.

Thus, in the general case, an iterative scheme like MCMC would be needed to sample from the posterior under our model, and such a scheme would be slow owing to the complexity of evaluating the target density.

Instead, we provide an algorithm for efficiently sampling entire paths from *p*(*X*_1:*N*_|*Y*_1:*N*_). This idea is due to Fearnhead (2006), with slight modifications to accommodate our model’s dependence between segment length and height.

Let *R*_*t*_ denote the event that a new IBD segment begins at position *t*, let 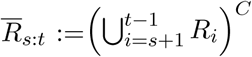 denote the event that there is *not* a recombination event between positions *s* and *t* (exclusive), and set 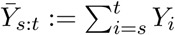. The joint likelihood of the data *Y*_*s*:*t*_ and the event that an IBD segment starts at position *s* and extends Δ = *t* −*s* + 1 positions before terminating at position *t* is

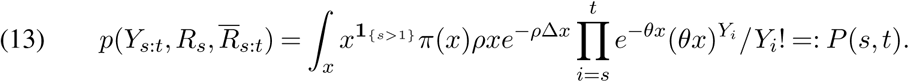

(A special case for *s* = 1 is necessary because the initial segment height is sampled from the stationary distribution *π*, while successive segments heights are distributed according to *<u>π</u>*; cf. equations 2 and 7).

For the last segment, we know only that it extended past position *N*, so make the special definition

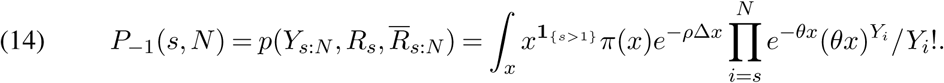

This algorithm can be used whenever (13) can be efficiently evaluated, in particular when *π*(*t*) is a gamma mixture. For example, if *π*(*x*) = *e*^−*x*^ then

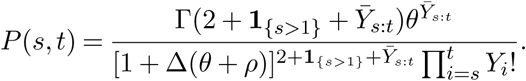

Defining *Q*(*t*) = *p*(*Y*_*t*:*N*_ |*R*_*t*_) and integrating over the location *s* where the segment originating at position *t* terminates, we have (Fearnhead, 2006, Theorem 1)

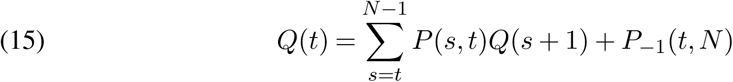

which can be solved by dynamic programming starting from *t* = *N* in *O*(*N* ^2^) time. As in the preceding section, when *t* −*s* is large, *P* (*s, t*) tends to be extremely small, so the summation in (15) can be truncated without loss of accuracy to obtain an algorithm which is effectively linear in *N*.

To sample the next recombination breakpoint *τ*′ from the posterior given that the previous breakpoint occurred at location *τ*, note that

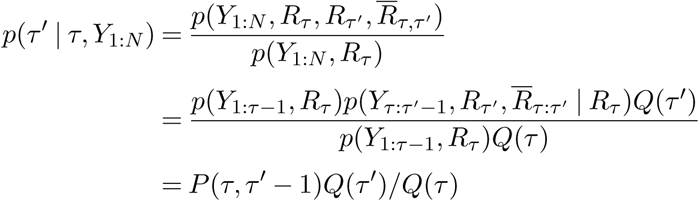

for *τ*′ = *τ* + 1, …, *N* − 1, with the remaining probability mass placed on the event that there are no more changepoints. If sampling the first changepoint we set *τ* = 1.

### 3.3. Exact frequentist inference

To complement the Bayesian results in the preceding section, we also derive an efficient frequentist method for inferring the *maximum a posteriori* (MAP) hidden state path,

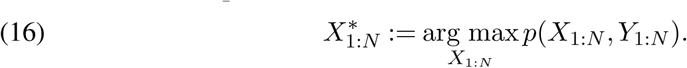

When *X*_1_, …, *X*_*N*_ ∈ 𝒳 have discrete support, |𝒳 | = *M*, the MAP path can be found in *O*(*NM* ^2^) time using the Viterbi algorithm (Bishop, 2006), and in some cases in *O*(*NM*) time by exploiting the special structure of the SMC (Harris et al., 2014; Palamara et al., 2018). Our goal is to solve the the optimization problem (16) in *O*(*N*) when 𝒳= ℝ_*>*0_.

To accomplish this, start by defining the recursive sequence of functions

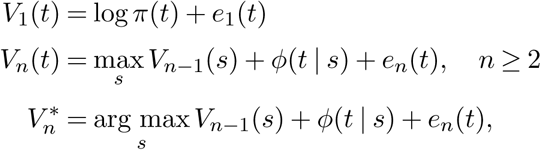

where *e*_*i*_(*t*) = log *p*(*Y*_*i*_ | *X*_*i*_ = *t*), and

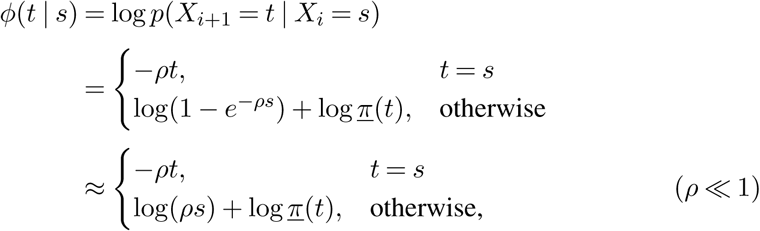

This is the usual Viterbi dynamic program, but defined over a continuous instead of discrete domain. By standard arguments (Bishop, 2006, §13.2.5), we have

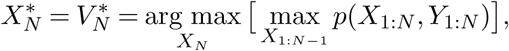

and the full path 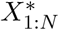 can be recovered by backtracing using the vector 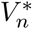.

Thus, if we could calculate *V*_*n*_(*t*) then the optimization problem (16) would be solved. In general, it is not obvious how to accomplish this, since *V*_*n*_(*t*) is a function, i.e., an infinite-dimensional object which cannot be represented by a computer program. However, our next theorem shows that, in fact, each *V*_*n*_(*t*) has a finite-dimensional representation.

#### Definition 1.

Let 𝒱_*K*_ be the space of all functions *f* : [0, ∞) → ℝ which can be piecewise defined by *K* functions of the form *t* ⟼ *at* + *b* log *t* + *c*. That is, *f* ∈ 𝒱_*K*_ if and only if there exists there exists an integer *K*, a vector ***τ*** ∈ ℝ^*K*+1^ satisfying

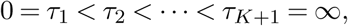

and vectors **a, b, c** ∈ ℝ^*K*^ such that

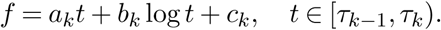

#### Theorem 2.

*For each n* = 1, …, *N, there exists K*_*n*_ *<* ∞ *such that* 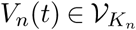.

To prove the result we need a few lemmas. We omit the trivial proofs of the first two.

#### Lemma 2.

𝒱_*K*_ *contains all piecewise constant and piecewise linear functions with K pieces. For all i, V*_*i*_ ⊂ *V*_*i*+1_. *If c* ∈ ℝ *and f* ∈ 𝒱_*i*_, *g* ∈ 𝒱_*j*_, *then cf* ∈ 𝒱_*i*_, *f* + *g* ∈ 𝒱_*i*+*j*_ *and* max{*f, g*} ∈ 𝒱_*i*+*j*_.

#### Lemma 3.

*Let e*_*n*_(*t*) := log *p*(*Y*_*n*_ | *X*_*n*_ = *t*). *Then*

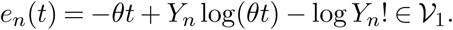

#### Lemma 4.

*Suppose that N*_*e*_(*t*) ∈ 𝒱_*K*_ *is piecewise constant. Then* log *<u>π</u>* (*t*) ∈ 𝒱_*K*_.

#### Proof.

If *N*_*e*_(*t*) is piecewise constant then so too is log *η*(*t*) = −log *N*_*e*_(*t*). Also, 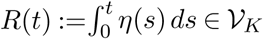 is piecewise linear on the same set of breakpoints. Hence,

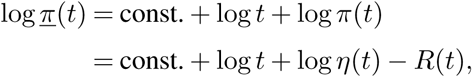

and the result follows from Lemma 2.□

#### Proof of Theorem 2.

By induction on *n*. For *n* = 1,

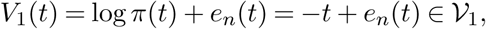

as claimed. For the inductive step, we have

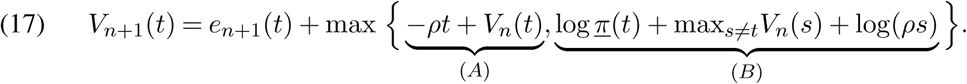

By the induction hypothesis and Lemmas 2-4, both (*A*) and (*B*) are in 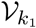 for some *k*_1_. Then, another application of the lemmas shows that in fact the entire right-hand side of (17) is in 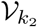 for some (possibly larger) *k*_2_.

#### Remark.

The proof shows that in order to efficiently compute *V*_*n*_(*t*) we need to be able to take the pointwise maximum between any two functions in 𝒱_*K*_. We provide an *O*(*K*) procedure for doing this in Appendix B.

Our next result shows that the structure of *V*_*n*_(*t*) essentially mirrors that of the posterior distribution *p*(*X*_*n*_|*Y*_1:*n*_) (cf. Section 4.2 and Appendix A). Each piece of *V*_*n*_(*t*) comprises an interval *I* ⊂ ℝ where, conditional on the TMRCA at position *n* being *t* ∈ *I*, the most probable recombination event occurred a certain number of positions ago. In the statement and proof of the theorem, we use double brackets, ⟦·⟧, to refer to individual entries of subscripted vectors; cf. Appendix A.

#### Theorem 3.

*For each V*_*n*_(*t*), *with breakpoints* 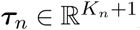, *there exists vectors* 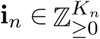 *and* 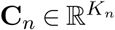 *such that, for t* ∈ [***τ***_*n*_⟦*k*⟧, ***τ***_*n*_⟦*k* + 1⟧),

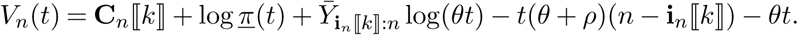

*Hence, up to the constant* **C**_*n*_ ⟦*k*⟧, *V*_*n*_ (*t*) *equals the log-likelihood of* 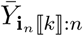 *given that the most recent recombination event occurred at position* **i**_*n*_ ⟦*k*⟧ *and* 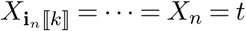.

#### Proof.

In view of equation (17), note that for fixed *t*, we can unwind the recursion *V*_*n*+1_(*t*) = *e*_*n*+1_(*t*) − *θt* + *V*_*n*_(*t*) until we reach an index *i* where (*A*) *<* (*B*). By continuity, this index is the same for all *t* [***τ***_*n*_ ⟦*k*⟧, ***τ***_*n*_ ⟦*k* + 1⟧). Denote the vector of such indices associated with each interval by **i**_*n*_, and let

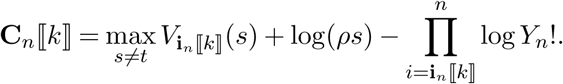

Then

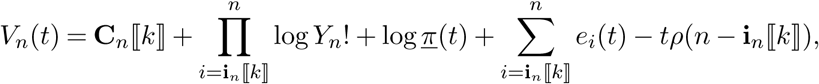

so the claim follows by using Lemma 3 to expand the sum.□

#### 3.3.1. Linear running time

Theorem 2 paves the way for inference by proving that each *V*_*n*_(*t*) can be finitely represented. However, it leaves open the possibility that the dimension *K*_*n*_ of this representation increases with *n*. This would imply that the running time of our algorithm increases faster than linearly in the sequence length *N*, an impediment to real-world applications.

Surprisingly, this does not happen: as we show empirically in Section 4.3, the recursion (17) is “self-pruning” in the sense that term (*B*) of that equation frequently dominates (*A*) over entire intervals, meaning that those terms can be dropped. This makes intuitive sense since (*B*) corresponds to the event that a recombination occurred between positions *n* and *n* + 1, and this will be the most likely explanation for extreme values of *t*.

Thus, we find that the average number of intervals tracked by our algorithm is bounded by a moderate constant, implying that the expected running time of this method is linear in the sequence length *N*. This result agrees with recent findings in the changepoint detection literature, where pruned dynamic programming has been used to derive methods whose average complexity grows linearly in the amount of data (Killick, Fearnhead and Eckley, 2012; Johnson, 2013; Maidstone et al., 2017).

#### 3.3.2. Generalization to non-MAP paths

It will be seen in Section 4.2 that the MAP path is rather different from a “typical” path sampled from the posterior distribution: the former tends to oversmooth, missing many recombination breakpoints, whereas the posterior mode is generally centered over the truth (Figure S5). This behavior occurs in hidden Markov models more generally, and can be understood in terms of decision theory (Yau and Holmes, 2013; Lember and Koloydenko, 2014; Kuljus and Lember, 2016). The MAP path 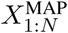 solves the optimization problem

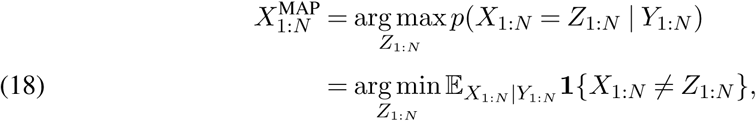

so the Viterbi algorithm can be interpreted as minimizing risk with respect to the loss function *ℓ*_MAP_(*x, y*) = **1** {*x* ≠ *y* }, where *x, y* are paths, and *x* = *y* if they are equal at every position. This loss function is “global” in that paths incur equal loss irrespective of whether they mismatch the true path at one position or all of them; there is no benefit to improving the match at a particular position.

On the opposite end of the spectrum, the pointwise posterior mode

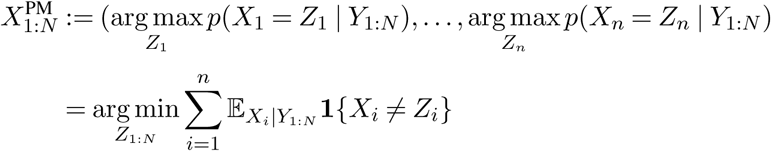

is “local”, placing no emphasis on paths that are continuous from one position to the next. Indeed, from Theorem 4 and Appendix A, we can see that arg max *p*(*X*_*i*_ | *Y*_1:*N*_) ≠ arg max *p*(*X*_*i*+1_ | *Y*_1:*N*_) almost surely for all *i*, so that 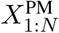 has a changepoint at every position and thus vanishingly small prior probability for large *N*.

For ordinary HMMs, it is possible to algorithmically interpolate between these these two extremes, resulting in paths that achieve better pointwise accuracy than 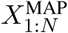 and higher prior likelihood than 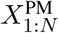 (Yau and Holmes, 2013; Lember and Koloydenko, 2014). However, these algorithms assume a discrete state space, and it is unclear whether they can be extended to our setting. Instead, we propose a simple modification of our method which has a straightforward interpretation as penalized changepoint detection.

To build the connection, note that we can write the optimization in (16) equivalently by representing *X*_1:*N*_ by the locations and heights of each segment, ***τ***, **x** ∈ ℝ^*K*^, such that

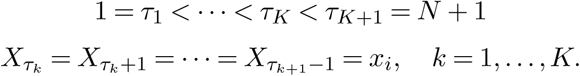

Then, we can rewrite the complete likelihood as

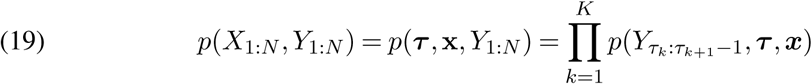

where

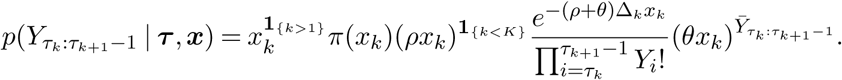

and Δ_*k*_ = *τ*_*k*+1_−*τ*_*k*_. Under the renewal approximation, for fixed ***τ***, (19) separates into a series of simpler one-dimensional optimization problems:

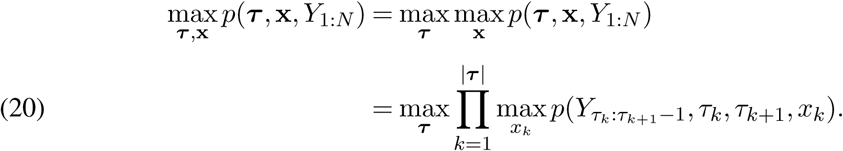

where we abused notation to write |***τ***| for the dimension of (i.e. the number of changepoints in) ***τ***. Taking the log of equation (20), we have that the MAP path equivalently solves

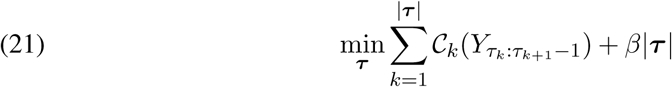

where we defined *β* = −log *ρ* and

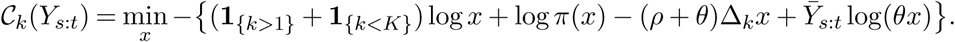

Hence, *β* penalizes segmentations with many changepoints. Above we showed that with *β* = − log *ρ*, the optimum of (21) is exactly 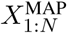, which is also optimal for (18). Other settings of *β* result in paths which are suboptimal with respect to this objective, but potentially superior by other metrics. In particular, we observed that by setting *β* lower than log *ρ*, thus encouraging the algorithm to find paths with more changepoints than the MAP path, the paths are pointwise superior to 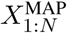 in the sense of the preceding paragraph.

### 3.4. Extension to larger sample sizes

The preceding sections focused on inferring the sequence of TMRCAs in a pair of sampled chromosomes. In modern applications where hundreds or thousands of samples have been collected, methods that can analyze larger sample sizes are both useful and desirable.

To generalize our inference problem to larger sample sizes, we recast it as follows: given a “focal” chromosome *f* and a “panel” 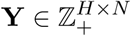, where *Y*_*hn*_ is the number of pairwise differences between chromosomes *f* and *h* at locus *n*, infer the sequence

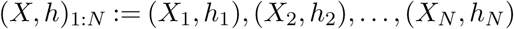

of genealogical nearest neighbors (GNNs) to *f*. In other words, at each position *i*, find the panel entry *h*_*i*_ ∈ {1, …, *H*} and corresponding TMRCA *X*_*i*_ for the chromosome most closely related to *f*. Note that there may be more than one GNN at a given site, so the sequence is not necessarily unique.

For *H* = 1 this problem reduces to finding *p*(*X*_1:*N*_|*Y*_1:*N*_), or its maximizer 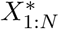, as outlined above. This suggests that we consider the likelihood *p*((*X, h*)_1:*N*_ |**Y**) in the general case. Unfortunately, evaluating this likelihood is significantly harder when *H >* 1. This is due to the fact that, except when *H* = 1, the sequence (*X, h*)_1:*N*_ does *not* uniquely determine the sequence of trees *T*_1_, …, *T*_*N*_ used to approximate the underlying ARG; we must integrate out the remaining uncertainty, which is quite difficult for the reasons described in Section 2.1. To circumvent this difficulty, we employ a so-called *trunk approximation*, which supposes that the ARG is a completely disconnected forest of *H* trunks extending infinitely far back into the past, in which case the sequence (*X, h*)_1:*N*_ is again in bijection with the (trivial) tree sequence *T*_1:*N*_. Although the trunk assumption is quite strong, it has proved useful in a variety of settings (Sheehan, Harris and Song, 2013; Spence et al., 2018; Steinrücken et al., 2019).

Modifying our methods to utilize the trunk approximation is straightforward and amounts to, essentially, replacing the coalescence measure *p*(*X* ∈ [*t, t* + *dt*)) = *π*(*t*) *dt* with the product measure *p*((*X, h*) ∈ ([*t, t* + *dt*), *i*)) = *π*(*Ht*) *dt* in all of our formulas. (Note that this measure is properly normalized.) In other words, coalescence occurs with each haplotype at rate 1, and conditional on coalescence, it occurs uniformly onto each haplotype.

## 4. Results

In this section we compare our method to existing ones, benchmark its speed and accuracy, and conclude with some applications.

### 4.1. Insensitivity of the posterior to the prior

As described in the introduction, the driving hypothesis of this work is that posterior inferences for the haplotype decoding problem are relatively insensitive to the choice of prior. In this section we investigate that hypothesis.

#### 4.1.1. Differences between different SMC models

We first studied whether different types of Markovian approximations to the spatial coalescent had a significant effect on the accuracy of posterior inferences. In particular, we initially compared the Markovian approximation of Hobolth and Jensen (2014) with the renewal approximation described above. SMC (McVean and Cardin, 2005) and SMC’ (Marjoram and Wall, 2006) are further approximations of the Markovian approximation to the tree building process, while the renewal process drops the dependence between neighboring tree heights altogether. Thus, the Markovian approximation and the renewal process can be viewed as the least and most approximative SMC methods, respectively.

To study the relationship between the posterior and prior, we compared the two methods under both constant population size and varying population size, as well as when the recombination rate is equal to the mutation rate and when it is lower. Under the constant size simulations, the effective population size was set to *N*_*e*_(*t*) = 20, 000 for all *t*. In the varying case,

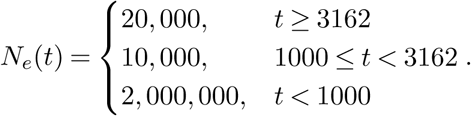

Taking all the combinations of the different population size histories and the recombination rate gives us a total of 4 scenarios. Scenarios 1 and 3 have constant population size, and scenarios 2 and 4 have the variable population size. Scenarios 1 and 2 have recombination rate *r* = 10^−9^, and scenarios 3 and 4 have recombination rate *r* = 1.4 × 10^−8^ per base-pair per generation. We bucket consecutive base pairs into groups of size *w* = 100 and assume that the recombinations occur between these groups. We discretized time into 32 epochs by selecting time points *t*_0_ = 0 *< t*_1_ *< < t*_32_ =∞ and setting epoch *I*_*ϵ*_ = [*t*_*ϵ*_, *t*_*ϵ*−1_). After setting the first time point as 0 and the final time point (*t*_32_) as ∞, we set *t*_1_, *t*_2_, …, *t*_31_ as the sequence of 31 evenly log-spaced numbers between 10 and 100, 000 including the endpoints.

In what follows we measure the accuracy of the discretized SMC posterior with respect to the true (simulated) TMRCA at each position. To do this, we assume that coalescence events occur at the expected time of coalescence given that coalescence occurred in that epoch. (See Appendix C for a precise description of our metrics.) To perform a fair comparison, even though we know how to solve the renewal model exactly, in this section we compare the time-discretized versions of it and the Markovian model.

For each scenario, we used `msprime` (Kelleher, Etheridge and McVean, 2016) to simulate *L* = 5 × 10^6^ base pairs of sequence data for 25 pairs of chromosomes, for a total of *L/w* = 5 × 10^4^ loci. The sequences were simulated with a per generation mutation rate of *µ* = 1.4 × 10^−8^. Note that in scenarios 3 and 4, *µ* = *r*. We calculate the posterior probabilities for the Markovian and renewal approximation using their corresponding transition probabilities. To assess the accuracy of the two priors we measure error using both an absolute and relative scale. We define the absolute error as

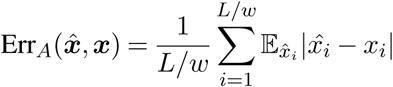

and the relative error as

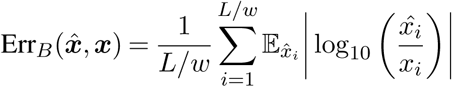

where *x*_*i*_ is the true TMRCA of the tree at position *i* and 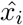 is time to coalescence distributed according to the posterior. The set of values that 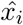 can take is 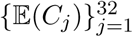 where *C*_*j*_ is the expected time to coalescence given that coalescence occurred in epoch *j* and is distributed according to the posterior at locus *i*.

Supplemental Figures S1 and S2 show the Viterbi path and the posterior heatmap for one run of each scenario of the simulation. Looking at all the panels of Figure S1, there is little difference in the Viterbi plot between the Markovian and renewal priors. Both priors produce a Viterbi path very similar to the true sequence of TMRCAs. When the recombination rate increases, the Viterbi path produced by the two priors fail to capture all the recombination events, but are still very similar in their outputs. In Figure S2 it is even more difficult to discern any meaningful difference in all scenarios between the two priors. This is especially the case in scenarios 1 and 2 where the recombination rate is lower.

Confirming these qualitative observations, Table 1 shows the average absolute error for the two priors over the 25 simulations. In terms of absolute error, the renewal prior does as well as the more correct Markov prior. In fact, the renewal prior outperforms the Markov prior under scenarios 3 and 4, the scenarios with higher recombination rate. Table S1 shows that the Markov approximation slightly is slightly better in relative error. However, in general the differences are minor, and both the tables confirm our hypothesis that the posterior is fairly insensitive to the choice of prior.

**Table 1.** Mean absolute error (Err_A_) over 25 runs under each scenario. Standard error in parenthesis.

| Scenario | 1 | 2 | 3 | 4 |
| --- | --- | --- | --- | --- |
| Markov | 6075.50 (215.59) | 5187.24 (187.27) | 12037.25 (298.71) | 12112.81 (106.44) |
| Renewal | 6068.04 (209.11) | 5187.49 (184.34) | 11476.92 (282.36) | 11571.28 (97.99) |

To further understand the difference between the priors, we stratified this analysis by quartiles of the true TMRCA. We denote the minimum and maximum TMRCAs as *q*_0_ and *q*_4_, and the first, second, and third quartiles as *q*_1_, *q*_2_, and *q*_3_. We then recalculate the absolute error in quarter *j* as

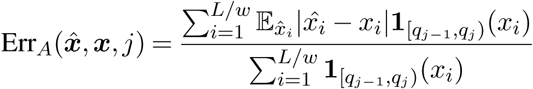

with relative error defined similarly. Due to the length bias of IBD tracts, the number of loci in quarter *j* will be smaller than the number of loci in quarter *j* − 1. The number of loci in each quarter under the various scenarios is displayed in Table S3.

Tables 2 contains the mean absolute error over the 25 simulations after stratification. Under scenarios 1 and 2 where the recombination rate is lower, again we see virtually no difference between the two priors across all quarters. Under scenarios 3 and 4 where the recombination rate is higher, we see that in the first and second quarters, the renewal prior outperforms the Markov approximation by a large margin. The results are reversed in the third and fourth quarters where the Markov approximation is more accurate than the renewal prior. This trend is mostly mirrored in Table S2 with the mean relative errors. The renewal prior does just slightly worse than the Markov prior under scenarios 1 and 2 across all quarters. Under scenarios 3 and 4 as the underlying true TMRCA increases, so too does the difference in Err_*B*_. The large difference in quarter 4 is expected as under the Markov prior, the distribution of tree height of the current segment conditioned on the tree height of the previous segment, *q*(*t*|*s*) is approximately uniform in *t* for large *s*; i.e. *q*(*t*|*s*) ≈ 1*/s* when *s* ≫ *t*. In contrast, the distribution under the renewal prior *π*(*t*) = *e*^−*t*^ is more dense for smaller values of *t*.

**Table 2.** Mean absolute error (Err_A_) over 25 runs under each scenario stratified by quartile. Standard error in parenthesis.

|  | Scenario | 1 | 2 | 3 | 4 |
| --- | --- | --- | --- | --- | --- |
| Markov | Q1 | 2840.79 (124.42) | 2219.61 (101.38) | 6820.01 (245.65) | 6672.23 (88.84) |
| Renewal | Q1 | 2887.14 (129.53) | 2273.91 (100.00) | 5292.75 (180.86) | 5256.89 (55.12) |
| Markov | Q2 | 6524.06 (131.61) | 6106.74 (122.89) | 13346.56 (35.50) | 13465.23 (57.68) |
| Renewal | Q2 | 6604.09 (117.57) | 6123.72 (107.90) | 11463.27 (38.05) | 11665.01 (35.60) |
| Markov | Q3 | 10543.82 (239.47) | 9716.04 (219.67) | 18831.10 (64.05) | 18957.97 (67.16) |
| Renewal | Q3 | 10454.31 (249.99) | 9528.27 (225.84) | 19538.07 (72.65) | 19682.76 (71.76) |
| Markov | Q4 | 17242.02 (460.92) | 16362.35 (471.86) | 33016.17 (212.12) | 33526.33 (226.10) |
| Renewal | Q4 | 16959.09 (505.93) | 16341.50 (601.40) | 40181.58 (263.39) | 40141.12 (279.70) |

In general, outside of the large difference between the methods in quarter 4, the two approximations are comparable, with neither one clearly dominating the other. When the underlying true TMRCA is smaller, Err_*A*_ is the better measure of accuracy, so despite the Markov approximation outperforming the renewal prior in all quarters in terms of Err_*B*_, the renewal prior actually outperforms the Markov approximation in quarters 1 and 2. We conclude from these results that our choice of prior is justified.

#### 4.1.2. Effect of the demographic prior

Next, we studied the extent to which the demographic prior *π*(*t*) affects the resulting estimates. We simulated data under three different demographic models and then measured the resulting accuracy of the posterior when each model was used as a prior to infer TMRCAs on data generated from the other models. The standard library for population genetic simulation models, `stdpopsim` (Adrion et al., 2019), provides a demographic model of the human population in Africa available as `Africa_1T12` and a difficult demography known in the literature as the zig-zag model (Schiffels and Durbin, 2014) available as `Zigzag_1S14`. This is a pathological model of repeated exponential expansions and contractions, and is designed to benchmark various demographic inference procedures.

In addition to these two models, we use a model with a constant population size of 2× 10^4^. We modeled the two non-constant population size history using a piecewise constant function of 64 segments instead of a continuous function. The three models are plotted in Supplemental Figure S3. The set of time breakpoints used to approximate the size history is also the same set of points we used to discretize time into epochs. Here we discretized time into 64 epochs setting *t*_0_ = 0, *t*_64_ =∞, and the sequence *t*_1_ *<* …*< t*_63_ as the sequence 63 evenly log-spaced numbers between 10 and 10^6^ including the endpoints.

We then simulated 25 pairs of chromosomes for each model with `msprime` using the human chromosome 20 model with the default flat recombination and mutation maps in conjunction with the demographic models. The per generation per base pair mutation rate and recombination rate for chromosome 20 given by `stdpopsim` are *µ* = 1.29 × 10^−8^ and *r* = 1.718 × 10^−8^ respectively. After simulating the data, for each pair of chromosomes generated under each of the models, we used each demographic size history as a demographic prior to calculate the posterior distribution of the TMRCA using the renewal approximation.

We display the posterior of one pair of chromosomes for all 9 pairs of demographies used as data generation and demographic priors in Figure S4. The plots show that regardless of which demographic prior was used, the resulting posteriors all had the same shape. There does seem to be a slight difference between the zig-zag demography and the other two demographies, in that the zig-zag posterior is generally more diffuse.

We use the same measures of accuracy as in the previous simulation, Err_*A*_ and Err_*B*_, to quantify how well the demographic priors perform against one another. Table 3 shows that in terms of mean absolute error, all three demographic models perform similarly when used as prior, regardless of which one of them in fact generated the data. Given the vast difference between the three demographic models (Figure S3), if the posterior were sensitive to the demographic model we would expect each column in the table to be quite different from one another. However, this is clearly not the case; using the correct prior results in an average improvement of a few percent in most cases.

**Table 3.** Mean absolute error (Err_A_) over 25 runs under each scenario. Standard error in parenthesis.

| Simulation Model | Prior Used |  |  |
| --- | --- | --- | --- |
|  | Africa | Zigzag | Constant |
| Africa | 10144.40 (19.76) | 10359.75 (20.84) | 10663.13 (20.19) |
| Zigzag | 5507.79 (132.03) | 4962.75 (120.63) | 5700.60 (137.21) |
| Constant | 11584.24 (259.28) | 11764.86 (263.18) | 11898.49 (266.09) |

Relative error measurements (Table S4) tell a similar story. The Africa and zig-zag demographies perform the best when used as a prior when they are also used to generate the data. In addition, the zig-zag demography performs somewhat worse than the other two demographies when the data is generated with the other demographies. While this does suggest that the posterior is somewhat sensitive to the prior, the zig-zag demography is an unrealistic model that would not be used in practice. In particular, it assumes that the population underwent repeated bottlenecks, causing estimated TMRCA to be low in general, and resulting in low absolute error rates in Table 3 and inflated relative error rates in Table S4. In conclusion, our results suggest that, as long as the chosen prior is not pathological, its effect on inference will be limited.

### 4.2. Comparison of Bayesian and frequentist inferences

In Section 3 we derived methods for inferring tree heights. Here we compare the Bayesian method where we sample from the posterior and the frequentist method where we take the MAP path. We apply these two methods to the same simulated data from Section 4.1.1. For the Bayesian method we sample 200 paths from the posterior and take the median to compare against the MAP path.

Figure S5 shows the results of running the two methods on one set of simulated chromosomes under each scenario. The top two panels of the figure show that when the recombination rate is an order of magnitude lower than the mutation rate, both methods give a faithful approximation of the true sequence of TMRCAs. However, the bottom two panels where the recombination rate is larger displays the key difference between the two methods: the MAP path fails to detect many recombination events, whereas the posterior median is an average over many paths so it can detect recombination events that the MAP path cannot.

We use the same measures of absolute and relative we used in the previous sections. For this simulation, we look at the error at each position so *L/w* = *L*. The results in Tables 4 and S5 show that the posterior median dominates the MAP path. Again, since the MAP path is the most likely single path whereas in the Bayesian method we take the pointwise median of many paths, the MAP path has inferior pointwise accuracy. This result is expected, but it should be noted that when compared to Tables 1 and S1, the MAP path performs similarly to, and the Bayesian method greatly outperforms, the posterior decoding of the discretized SMC models.

**Table 4.** Mean absolute error (Err_A_) over 25 runs under each scenario. Standard error in parenthesis.

| Scenario | 1 | 2 | 3 | 4 |
| --- | --- | --- | --- | --- |
| MAP | 6021.78 (234.36) | 5040.54 (183.42) | 11916.00 (293.34) | 12060.40 (122.81) |
| Bayesian | 4422.97 (173.91) | 3845.58 (139.81) | 8636.69 (197.69) | 8531.71 (78.97) |

### 4.3. Empirical running time

In Sections 3.2 and 3.3.1 we suggested that by pruning the state space of our methods in certain ways, their running time is effectively linear in the number of decoded positions. A rigorous proof of this fact is difficult, and beyond the scope of this article. We settled for verifying that the claim holds in simulations. We benchmarked our methods on simulated sequences of length *L* = 10^6^ to *L* = 10^8^. For each length, we simulated 10 pairs of chromosomes. Figure S6 confirms that there is a linear relationship between chromosome length and running time for both the Bayesian sampler method and the MAP decoder. Note that, if decoding against a larger panel of chromosomes (cf. Section 3.4), the amount of work performed by our algorithms scales linearly in the panel size *H*. Therefore, the empirical running time of our methods is *O*(*HL*). This matches the running time of the most efficient existing methods for decoding the SMC (Harris et al., 2014; Palamara et al., 2018).

### 4.4. Applications

We tested our method on the two most common real-world applications of the sequentially Markov coalescent.

#### 4.4.1. Exact PSMC

The pairwise sequentially Markov coalescent (PSMC; Li and Durbin, 2011) is a method for inferring historical population size (i.e., the function *N*_*e*_(*t*) defined in Section 2.2) using genetic variation data from a single diploid individual. Although in some settings PSMC has been superseded by more advanced methods which can analyze larger sample sizes (Schiffels and Durbin, 2014; Terhorst, Kamm and Song, 2017; Steinrücken et al., 2019), it remains very widely used in many areas of genetics, ecology and biology, because it is fairly robust, and does not require phased data, which can be difficult to obtain for species that have not been studied as intensively as humans.

As noted in Section 1, PSMC uses an HMM to infer a discretized sequence of genealogies. The discretization grid is a tuning parameter which is challenging to set properly—finer grids inflate both computation time and the variance of the resulting estimate, and for a fixed level of discretization, the optimal grid depends on the unknown quantity of interest *N*_*e*_(*t*). A poorly chosen discretization can have serious repercussions for inference (Parag and Pybus, 2019). It is preferable to dispense with this tuning parameter altogether, as our method enables us to do.

A second benefit of our approach is that it allows us to recast the problem in a more natural form. PSMC requires the user to fix a parametric function class for *N*_*e*_(*t*), also dependent on the aforementioned discretization, and performs parameter optimization via E-M. This process is slow and occasionally unstable. We will proceed differently, by establishing a connection to density estimation. Recalling equation (3), we see that inference of *N*_*e*_(*t*) is tantamount to estimating (the reciprocal of) *η*(*t*). In survival analysis, *η* is known as the *hazard rate function*, and a variety of methods have been developed to infer it (Wang, 2014). Thus, if we could somehow sample directly from *π*, then inference of *N*_*e*_(*t*) would reduce to a fairly well-understood problem.

While this is impossible in practice, the simulated results shown in the preceding sections inspire us to believe that samples drawn from the posterior *p*(*X*_1:*N*_|*Y*_1:*N*_) could serve the same purpose. Concretely, we suppose that a random sample *x*_1_, …, *x*_*k*_ drawn from the product measure

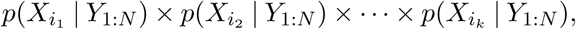

where the index sequence *i*_1_, …, *i*_*k*_ ∈ [*N*] is sufficiently separated to minimize correlations between the posteriors, is distributed as *k* i.i.d. samples from *π*. We then use a kernel-smoothed version of Nelson-Aalen estimator (Wang, 2014) in order to estimate 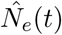.

We first compared the performance of our method with PSMC on simulated data. Figure 1 compares the results of running our method, which we call XSMC (e**X**act **SMC**), and PSMC on data simulated from three size history functions (plotted as dashed grey lines). We simulated a chromosome of length *L* = 5 × 10^7^ base pairs for 25 diploid individuals (total of 50 chromosomes), and then ran both methods on all 25 pairs. The plots show the pointwise median, with the interquartile range (distance between the 25th and 75th percentiles) plotted as an opaque band around the median. For the first two simulations we assumed that the mutation and recombination rates were equal, *µ* = *r* = 1.4 × 10^−8^ per base pair per generation. For reasons discussed below, we assumed in the third simulation that *r* = 10^−9^. Both methods were run with their default parameters and provided with the true ratio *r/µ* used to generate the data.

**Fig 1.**
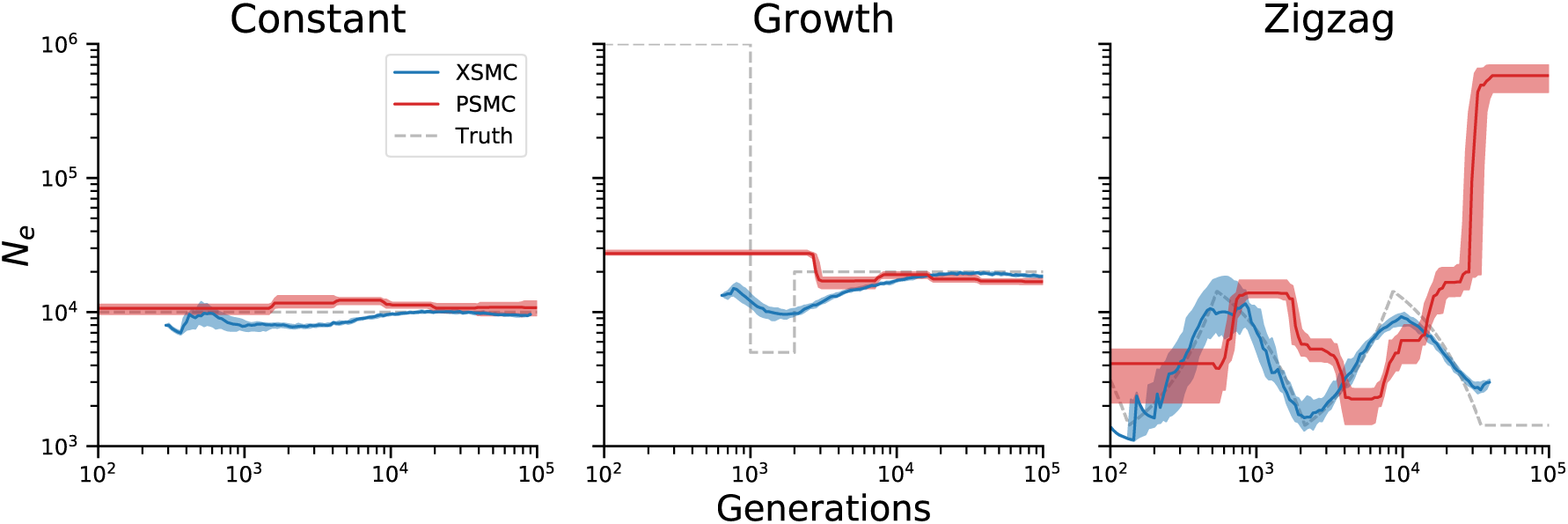
Comparison of PSMC and XSMC on various simulated size histories.

The left panel of the figure (“Constant”) depicts the most basic scenario, where the population size is unchanged over time. Both methods do an acceptable job, while exhibiting some bias. For PSMC, there is clear bias from the piecewise-constant model class it uses to perform estimation. XSMC has a slight downward bias in the recent past, but is otherwise centered over the true values *N*_*e*_ = 10^4^. Both methods appear slightly biased in the period 10^3^-10^4^ generations, though in opposite directions.

In the center panel (“Growth”), we simulated a cartoon model of recent expansion, in which the population experiences a brief bottleneck from 2,000–1,000 generations ago, before suddenly increasing in size by two hundredfold. This model is more difficult to correctly infer using only diploid data, because the large recent population size prevents samples from coalescing during this time, effectively depriving methods of the ability to learn size history in the recent past. Nevertheless, XSMC does an acceptable job of showing that the population experienced a dip followed by a sharp increase, though the estimates are oversmoothed. In contrast, PSMC estimates size history that is nearly flat, with no acknowledgement of the bottleneck. This result also illustrates another benefit of the nonparametric approach: XSMC only returns an answer where it actually observes data. Because no coalescence times were observed before ∼10^3^ generations when sampling from the posterior, our method does not plot anything outside of that region. This compares favorably with PSMC and related parametric methods (e.g., Schiffels and Durbin, 2014; Terhorst, Kamm and Song, 2017), which have to model *N*_*e*_(*t*) over all 0 ≤ *t <* ∞ in order to perform an analysis, even when the data contain no signal outside of a limited region.

Lastly, in the right-hand panel we examined the zig-zag demography mentioned previously in Section 4.1.2. We found that with the default setting *ρ* = *θ* used in the preceding two examples, neither XSMC nor PSMC produced good results on the zig-zag. We therefore lowered the rate of recombination to *r* = 10^−9^/bp/generation in order to create more linkage disequilibrium for the methods to exploit. Here, a fairly substantial difference emerges between the two methods. XSMC does a good job of inferring this difficult size history, with accurate results to almost 10^2^ generations in the past, and almost no discernible bias. In contrast, PSMC is very inaccurate, with a wildly overestimated ancestral population size, and an overall shape that differs substantially from the truth.

Encouraged by these results, we next turned to analyzing real data. We performed a simple analysis where we analyzed whole genome data from 20 individuals from each of the five superpopulations (African, European, East Asian, South Asian, and Admixed American) in the 1000 Genomes dataset (The 1000 Genomes Project Consortium, 2015). Results are shown in Figure 2. Broadly speaking, our method agrees with other recently-published estimates (Li and Durbin, 2011; Terhorst, Kamm and Song, 2017), and succeeds in capturing major recent events in human history such as an out-of-Africa event 100-200kya, a bottleneck experienced by non-African populations, and explosive recent growth beginning around 20kya. We suspect that these estimates could be improved somewhat with fine-tuning and the use of additional data, but we did not do this, the message being that our method has moderate data requirements and produces reasonable results with minimal user intervention. Finally, we note that our method is highly efficient: to analyze all 20 × 5 × (3 × 10^9^Mbp) ≈ 300Gbp of sequence data took approximately 40 minutes on a 12-core workstation. A single human genomes (all 22 autosomes) can be analyzed in about 30 seconds.

**Fig 2.**
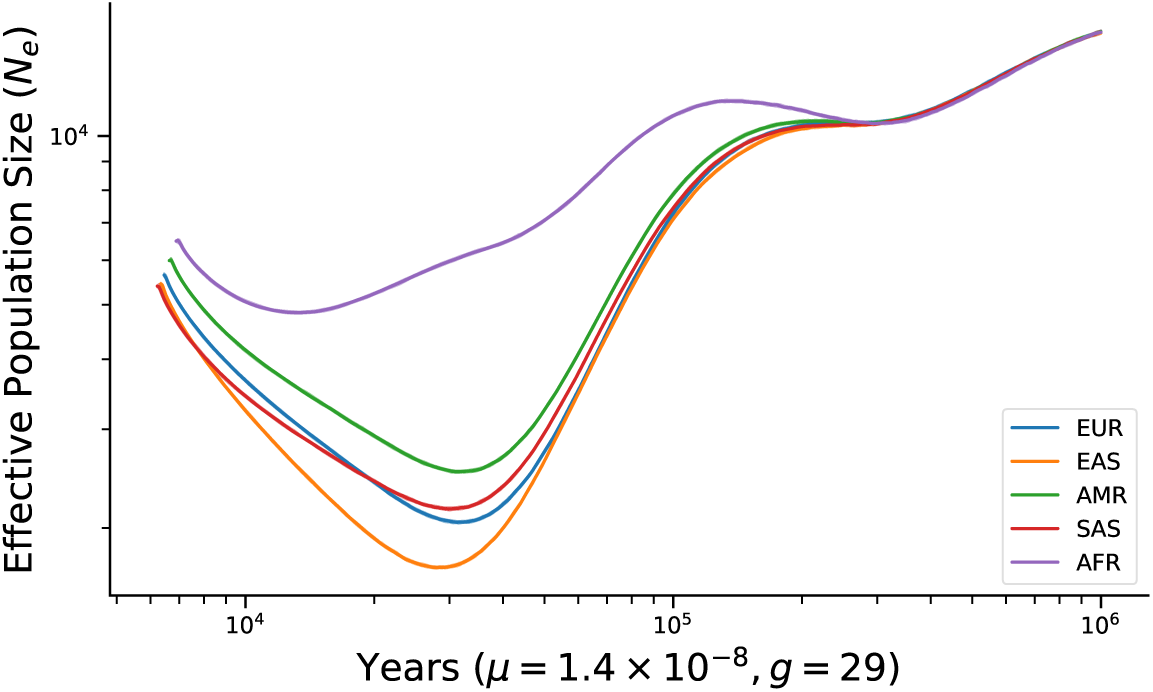
Result of fitting XSMC to 1000 Genomes data. For each superpopulation, 20 samples were chosen. Solid line denotes the median across all samples, and shaded bands denote the interquartile range.

#### 4.4.2. Phasing and Imputation

The Li and Stephens (2003) haplotype copying model (hereafter, LS) is an approximation to the conditional distribution of a “focal” haplotype (e.g., a chromosome) given a set of other “panel” haplotypes. It supposes that the focal haplotype copies with error from different members of panel, occasionally switching to a new template due to recombination. Genealogically, this can be interpreted as finding the local genealogical nearest neighbor (GNN) of the focal haplotype within the panel. LS has been used extensively in applications, for example phasing diploid genotype data into haplotypes (Stephens and Scheet, 2005) and imputing missing data (Li and Abecasis, 2006; Scheet and Stephens, 2006; Marchini et al., 2007; Howie, Donnelly and Marchini, 2009). The method’s undeniable success is actually somewhat surprising, since it assumes an extremely simple genealogical relationship between the focal and panel haplotypes which ignores time completely (Paul and Song, 2010; Paul, Steinrücken and Song, 2011). Hence, while we motivated XSMC as a fast and slightly more approximate SMC prior, it can also be seen as a more biologically faithful version of LS.

We wondered whether our method could be used to improve downstream phasing and imputation. Fully implementing a phasing or imputation pipeline is beyond the scope of this paper, so we settled for checking in simulations whether decoding results produced by XSMC were more genealogically accurate than those obtained using LS. We simulated data using realistic models of human chromosomes 10 and 13 (Adrion et al., 2019). We chose these two because chromosome 10 is estimated to have an average ratio of recombination to mutation slightly above 1 (*ρ/θ* = 1.07), while in chromosome 13 the ratio is slightly below 1 (*ρ/θ* = 0.87). The ratio of recombination to mutation affects the difficulty of phasing and imputation, with higher ratios leading to less linkage disequilibrium and thus less accurate results. We also explored the effects of varying the size of the haplotype panel. For each chromosome, we simulated 25 data sets with panels of size *H* = 2, 4, 10, 25, 100.

As a proxy for phasing and imputation accuracy, we studied which method identified a genealogical nearer neighbor on average. The GNN at a given position is defined to be any panel haplotype that shares the earliest common ancestor with the focal haplotype. In other words, any panel haplotype that has the smallest TMRCA with the focal haplotype is a GNN. (Note that there may be more than one GNN.) For purposes of accurate phasing and imputation, it is desirable to identify the GNN as closely as possible.

For each simulation we computed the Viterbi path from XSMC and LS, and studied the proximity of those paths to the true GNN at each segregating site. Table 5 shows the proportion of segregating sites where XSMC and LS both estimated the same haplotype to be the GNN. There is a high level of agreement, 80-90%, between the two methods for both small and large panel sizes. When the panel size is small (*H* = 2), there are few possible choices, and when the panel size is large (*H* = 100) the decoding consists mostly of long, recent stretches of IBD which are fairly easy to estimate. Disagreement is highest for intermediate values *H* = 4, 10, 25 where neither of these effects dominates. At sample size *H* = 10 the methods only agree at about half of segregating sites.

**Table 5.** Proportion of segregating sites where XSMC and the Li and Stephens’ agree on the GNN.

| Panel Size | 2 | 4 | 10 | 25 | 100 |
| --- | --- | --- | --- | --- | --- |
| Chromosome 10 | 0.9042 (0.0031) | 0.6653 (0.0047) | 0.5537 (0.0019) | 0.7738 (0.0019) | 0.8390 (0.0053) |
| Chromosome 13 | 0.9054 (0.0030) | 0.6664 (0.0051) | 0.5470 (0.0028) | 0.7787 (0.0019) | 0.8387 (0.0036) |

At the 10-50% of sites where the methods disagree, the results indicate a statistically significant gain for XSMC compared to LS. Table 6 shows that conditional on the two methods inferring different haplotypes as the GNN at that site, XSMC finds a genealogical nearer neighbor more often in all scenarios. This difference is most pronounced at the intermediate panel sizes, *H* = 4, 10, where XSMC selects a closer neighbor more than 80% of the time. The advantage of using XSMC increases, albeit slightly, in chromosome 13, indicating that our method may have an advantage when *ρ/θ* is smaller.

**Table 6.** Proportion of segregating sites that XSMC finds the more closely related haplotype than the Li and Stephens’ method conditional on the two methods inferring different haplotypes at that site.

| Panel Size | 2 | 4 | 10 | 25 | 100 |
| --- | --- | --- | --- | --- | --- |
| Chromosome 10 | 0.6860 (0.0018) | 0.8142 (0.0041) | 0.8351 (0.0015) | 0.6838 (0.0127) | 0.6528 (0.0058) |
| Chromosome 13 | 0.7160 (0.0013) | 0.8322 (0.0038) | 0.8495 (0.0023) | 0.7037 (0.0122) | 0.6766 (0.0112) |

## 5. Conclusion

In this article, we studied the sequentially Markov coalescent, a framework for approximating the likelihood of genetic data under various evolutionary models. We proposed a new inference method which supposes that the heights of neighboring identity-by-descent segments are independent. We showed that this led to decoding algorithms which are faster and have less bias than existing algorithms.

There are several possible extensions to our work. It is straightforward to extend our techniques to allow for position-specific rates of recombination and mutation, which could then be used to infer spatial or motif-specific variation in these processes.

Although we focused here on analyzing data from a single, panmictic population, we can also use posterior samples or MAP estimates to infer more complicated models of population structure. It is also possible to extend some of our techniques to other priors which model correlations between adjacent IBD segments. For the Viterbi decoder, we were able to implement a version of the algorithm in Section 3.3 which works for McVean and Cardin‘s original SMC model. This could be useful, for example, if analyzing data from a structured population, to the extent that adjacent segments of identity by descent are more likely to derive from members of the same subpopulation. However, the resulting procedure is much more complicated. The Viterbi function *V*_*n*_(*t*) no longer has the tractable form derived in Theorem 2. Consequently, we cannot use a simple method like the one in Appendix B to perform the pointwise maximization in equation (17). Instead, numerical optimization must be used instead, resulting in a slower algorithm. Similarly, on the Bayesian side, though there has been some work on posterior inference for correlated changepoint models (Fearnhead and Liu, 2011), these methods are slower and no longer sample exactly from the posterior.

Another interesting possibility is to use our method to estimate ancestral recombination graphs. Recently, there has been a resurgence of interest in inferring ARGs using large samples of cosmopolitan genomic data (Kelleher et al., 2019; Speidel et al., 2019). Although these represent an impressive breakthrough, they rely on heuristic estimation procedures that do not directly model the underlying genealogical process that generates ancestry. Our method provides a new possibility for ARG estimation, by iteratively “threading” each additional samples onto a sequence of estimated genealogies, but without the need to discretize those genealogies (Rasmussen et al., 2014). These and other extensions are the subjects of ongoing work.

## Data and code availability

All of the data analyzed in this paper are either simulated, or publicly available. A Python package implementing our method is available at https://terhorst.github.io/exact_smc. Code which reproduces all of the figures and tables in this manuscript is available at https://terhorst.github.io/exact_smc/paper.

## APPENDIX A FORWARD RECURSION CONSTANTS

In this section we derive the exact mixing weights and scale/shape parameters for the mixture representation proved in Theorem 1. Define *γ*(*x*; *a, b*) to be the PDF of the gamma distribution with mean *a/b* and variance *a/b*^2^,

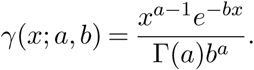

To conserve notation, in this section we use the following array-based conventions for vector expressions:

- Scalar functions operate on vectors in a component-wise manner. For example, if **x, y, z** ∈ ℝ^*k*^ then

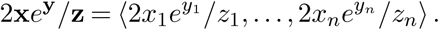
- In particular, for vectors ***α*, a, b** ∈ ℝ^*n*^,

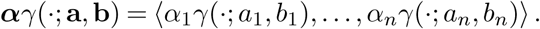
- A binary operation between a scalar and a vector “broadcasts” the scalar to the dimension of the vector. For example, 1 + **x** = ⟨1 + *x*_1_, …, 1 + *x*_*n*_⟩.
- To refer to individual entries of subscripted vectors, we will use the notation ⟦*i*⟧. A subvector (“slice”) of *x*_*n*_ of length *i* ≤ *n* is denoted **x**_*n*_ ⟦1 : *i*⟧ = ⟨ **x**_*n*_ ⟦1⟧, **x**_*n*_ ⟦2⟧, …, **x**_*n*_ ⟦*i*⟧ ⟩.
- The sum of all the entries of **x** is denoted 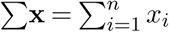

Additionally, we define the following function for later use:

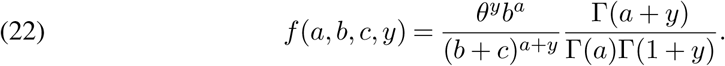

We prove the following theorem in the case where *π* is a gamma distribution. Extending the proof to gamma mixtures requires no new ideas, only notation; details are left to the reader.

### Theorem 4.

*Suppose that π*(*x*) = *γ*(*x*; *a*_0_, *b*_0_). *For each n* ∈ [*N*] *let* **a**_*n*_, **b**_*n*_ ∈ ℝ^*n*^ *be defined by*

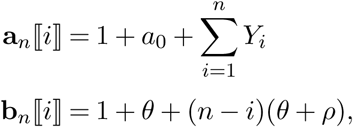

*and define* 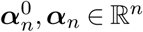, ***α***_*n*_ ∈ ℝ^*n*^ *and C*_*n*_ ∈ ℝ *by the recursions*

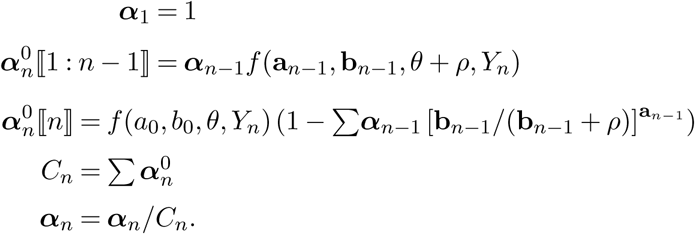

*Then*

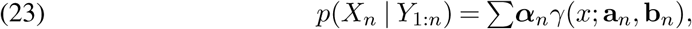

*and additionally C*_*n*_ = *p*(*Y*_*n*_ | *Y*_1:*n*−1_).

### Proof.

By induction on *n*. The base case *p*(*X*_1_ | *Y*_1_) follows from Fact 1 in the main text. For the general case, assume that *p*(*X*_*n*_ | *Y*_1:*n*_) has the form shown in (23). Then

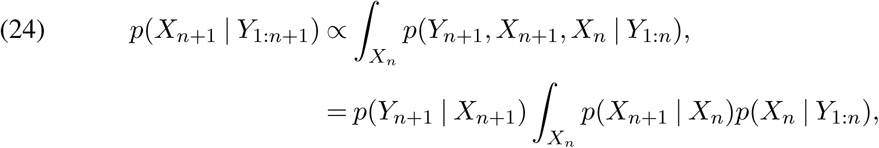

where the constant of proportionality *C*_*n*+1_ = *p*(*Y*_*n*+1_ |*Y*_1:*n*_) does not depend on *X*_*n*+1_. Using the transition rule (4), this implies

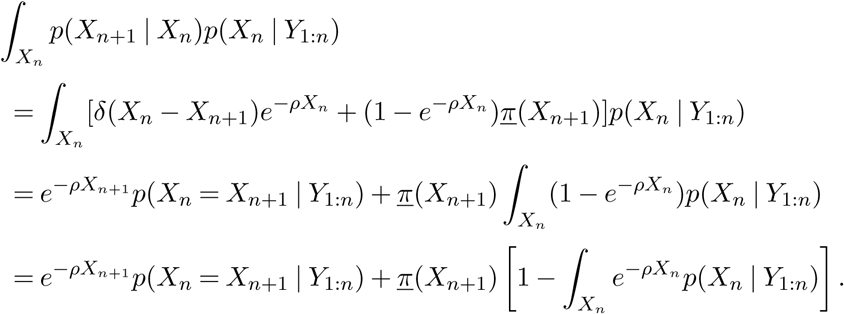

Now, by the inductive hypothesis and the identity

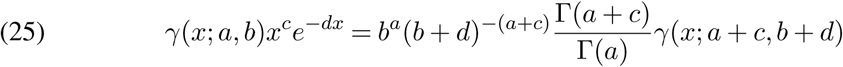

we obtain, for 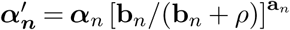

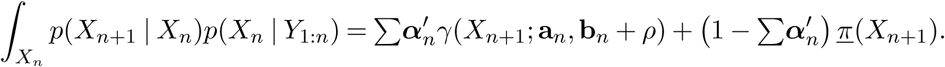

Multiplying through by

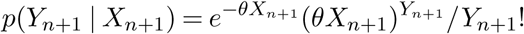

yields

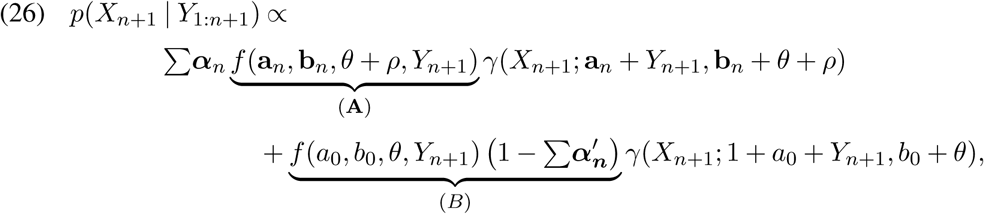

by (22) and (25). If we make the additional definitions

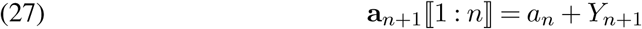

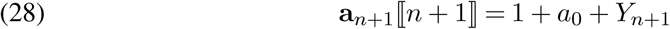

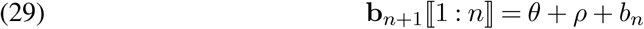

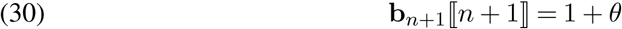

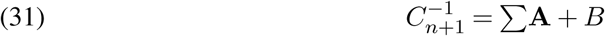

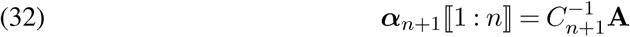

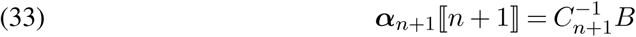

then (26) can be written as

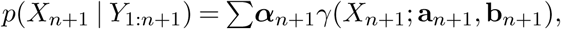

completing the proof. The recursive definition for ***α***_*n*+1_ follows from (32) and (33), and the representations for **a**_*n*+1_ and **b**_*n*+1_ follow from (27)–(30). Finally, note that *C*_*n*+1_ is precisely the constant of proportionality in (24) and therefore equals the conditional evidence *p*(*Y*_*n*+1_ | *Y*_1:*n*_).□

## APPENDIX B COMPUTING THE POINTWISE MAXIMUM IN V_*K*_

In this section we derive a procedure for finding the pointwise maximum max *f, g* when *f, g* ∈ 𝒱_*K*_. By enlarging *K* if necessary, we can without loss of generality assume that *f* and *g* are defined on the same set of breakpoints. Then it suffices to show how to find the zeros (if any) of the function *h* = *f* − *g* ∈ 𝒱_*K*_ over any given interval.

Accordingly, let *h*(*t*) = *at* + *b* log *t* + *c* for *τ* ∈ *I* := [*τ*_1_, *τ*_2_). By the change of variables −log *t* → *u* it is equivalent, and slightly simplifies the math, to find the zeros of *h*(*u*) = *ae*^−*u*^ − *bu* + *c* over an arbitrary interval *I*. Since interchanging the roles of *f* and *g* does not change the result, we may also assume that *a* ≥ 0, and if *a* = 0 then we may assume that *b* ≥ 0.

Let *w* = −*ae*^*c/b*^*/b*. If *w* ≥ 0 then *h*(*u*) has a single real root *u*_0_ = *W*_0_(*w*) − *c/b*, where *W*_0_(*x*) denotes the principal branch of the Lambert *W* function (DLMF, §4.13). If −1*/e* ≤ *w*_0_ *<* 0 then *h*(*u*) has two real roots, one at *u*_0_ and the other at *u*_1_ = *W*_−1_(*w*) − *c/b* where *W*_−1_ is the −1 branch of the Lambert *W* function.

We will use repeatedly the fact that a trivial solution exists whenever *h* can be shown to be globally decreasing, since:

- If *h*(*τ*_2_) ≥ 0 then the function is non-negative over *I*, so the maximum is *f*.
- If *h*(*τ*_1_) *<* 0 then the function is negative over *I*, so the maximum is *g*.
- Else the function has a single root *u*_0_ ∈ *I*, so the maximum is *f* on [*τ*_1_, *u*_0_) and *g* on [*u*_0_, *τ*_2_).

To find the zeros of *h*(*u*), we proceed by cases:

- If *b* = 0:
  − If *a* = 0 then *h* = *c*, so the maximum over *I* is either *f* or *g* depending on the sign of *c*.
  − Else (*a* ≥ 0, *b* = 0):
    * If *c* ≥ 0 then *h* = *f* − *g* ≥ 0 so the maximum over *I* is *f*.
    * Else, we have *h*′ = −*uae*^−*u*^ + *c <* 0 so the function is decreasing.
- If *a* = 0 then we assume that *b* ≥ 0. Then *h*′(*u*) = −*b* ≤ 0, so *h* is decreasing.
- Else (*a >* 0, *b* ≠ 0):
  − If *b >* 0 then *h*′(*u*) = *ae*^−*u*^ *b <* 0 so *h* is decreasing.
  − Else we have *h*^*′′*^(*u*) = *ae*^−*u*^ *>* 0 so the function is convex with a global minimum at *u*^∗^ = log(−*a/b*):
    * If *h*(*u*^∗^) *>* 0 then the function is non-negative so the maximum is *f*.
    * Otherwise, *h* is convex with

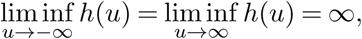

so it has two real roots *u*_0_ and *u*_1_. Without loss of generality assume *u*_0_ ≤ *u*_1_. There are 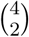 cases to consider depending on the ordering of *u*_0_, *u*_1_, *τ*_1_, *τ*_2_. For example, if *τ*_1_ *< u*_0_ *< u*_1_ *< τ*_2_ then *h* is positive on [*τ*_1_, *u*_0_), negative on [*u*_0_, *u*_1_) and positive on [*u*_1_, *τ*_2_), leading to a pointwise maximum function which takes on the values *f, g, f* on those three intervals. The other five cases are handled similarly, and we omit the details.

The running time of this procedure is *O*(1) assuming we can evaluate *W*_*n*_(*w*) in constant time. Thus, to find the pointwise maximum of *f* and *g* when both have are defined on *K* pieces takes *O*(*K*) time.

## APPENDIX C EXPECTED TIME TO COALESCENCE

In this section we describe how to calculate the expected time to coalescence which we use in the simulations in Sections 4.1.1 and 4.1.2. Suppose we have discretized time into the following set of *m* + 1 times points *t*_0_ = 0 *< t*_1_ *<* … *< t*_*m*_ = ∞. Precisely, the distribution of time to coalescence within epoch *I*_*ϵ*_ = [*t*_*ϵ*−1_, *t*_*ϵ*_) is

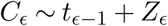

where *Z*_*ϵ*_ is a truncated exponential in the interval [0, *t*_*ϵ*_ − *t*_*ϵ*−1_) with parameter *η*_*ϵ*_ = 1*/N*_*e*_(*t*_*ϵ*−1_). The expectation of *Z*_*ϵ*_ is

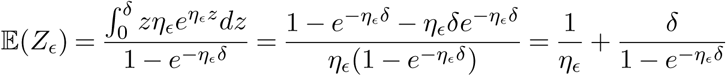

where *δ* = *t*_*ϵ*_ −*t*_*ϵ*−1_.

Finally, with some algebra we have that

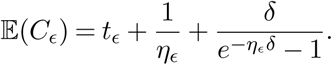

The final epoch *I*_*m*_ = [*t*_*m*−1_, *t*_*m*_) = [*t*_*m*−1_, ∞) is not bounded above, so the time to coalescence simply follows an exponential random variable with parameter *η*_*m*−1_ without trun-cation. Thus the expected time to coalescence is simply given by

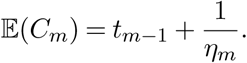

## SUPPLEMENTARY MATERIAL

### Additional figures and tables

**Fig S1.**
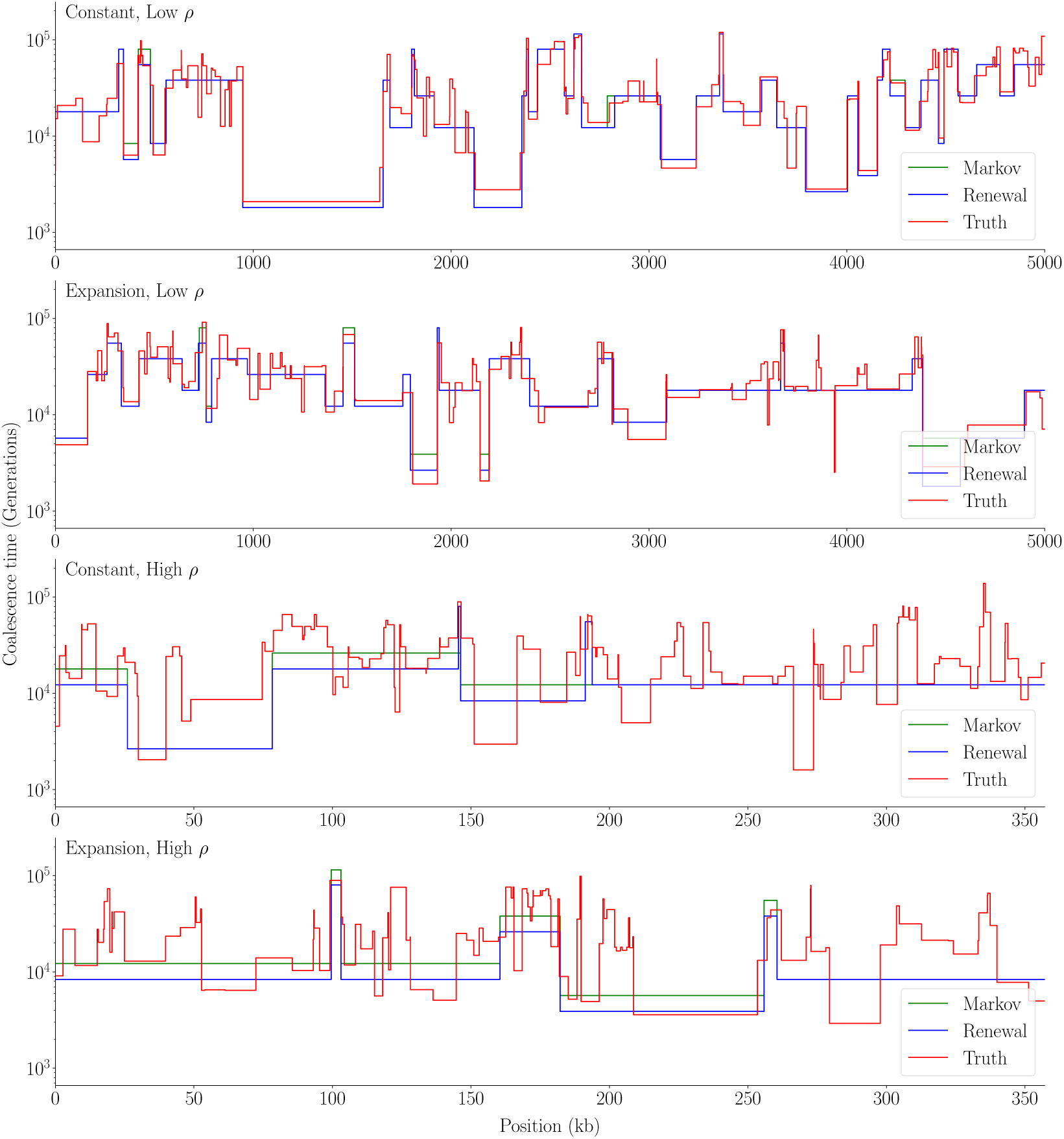
Comparison of Viterbi path between Markovian and renewal approximations.

**Table S1.** Mean relative error (Err_B_) over 25 runs under each scenario. Standard error in parenthesis.

| Scenario | 1 | 2 | 3 | 4 |
| --- | --- | --- | --- | --- |
| Markov | 0.1362 (0.0035) | 0.1281 (0.0024) | 0.3051 (0.0013) | 0.3006 (0.0014) |
| Renewal | 0.1437 (0.0026) | 0.1359 (0.0025) | 0.3451 (0.0046) | 0.3422 (0.0014) |

**Fig S2.**
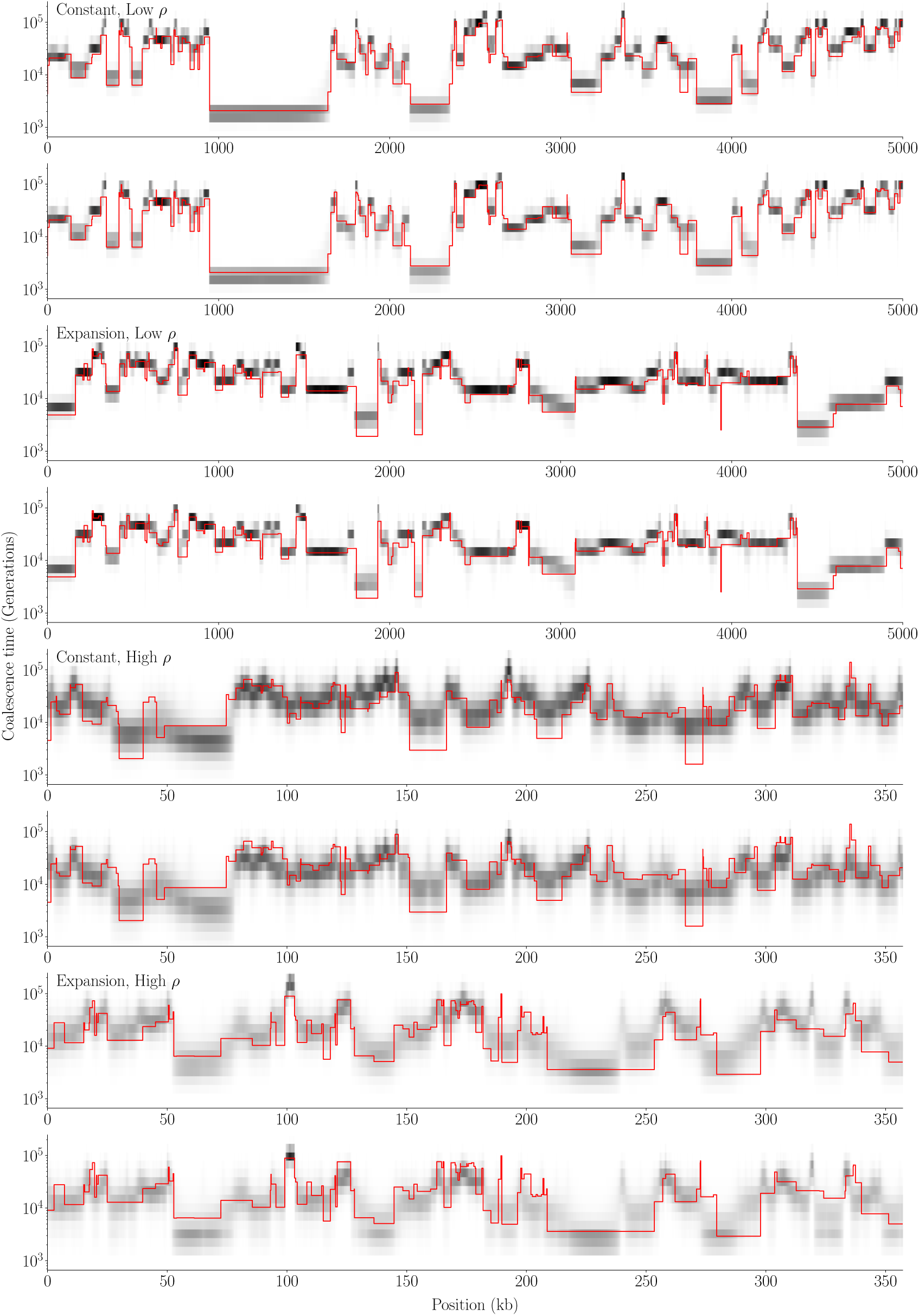
Comparison of posterior heatmap between Markovian and renewal approximations. The top panel in each group is the posterior given by the Markovian prior and the bottom panel is the posterior given by the renewal prior.

**Fig S3.**
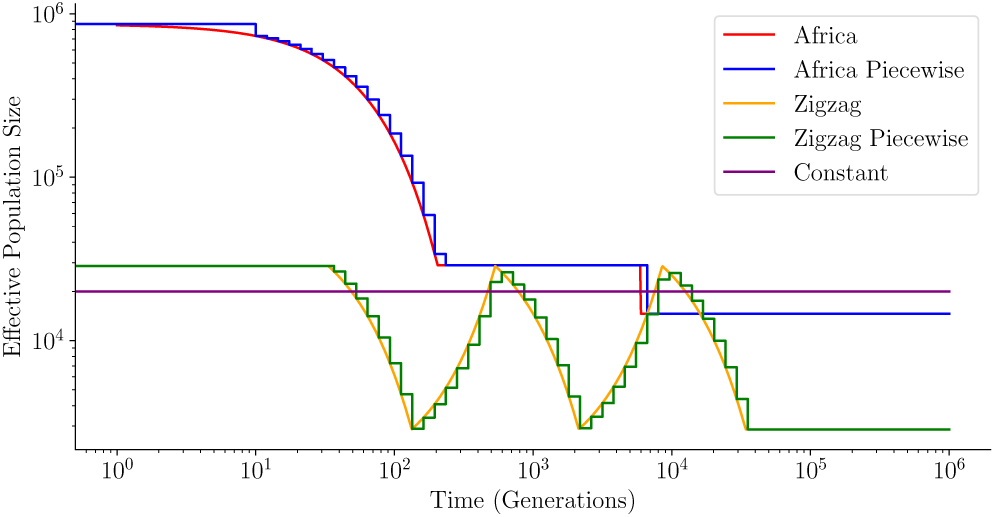
The population trajectory under the three models used in the simulation.

**Table S2.**
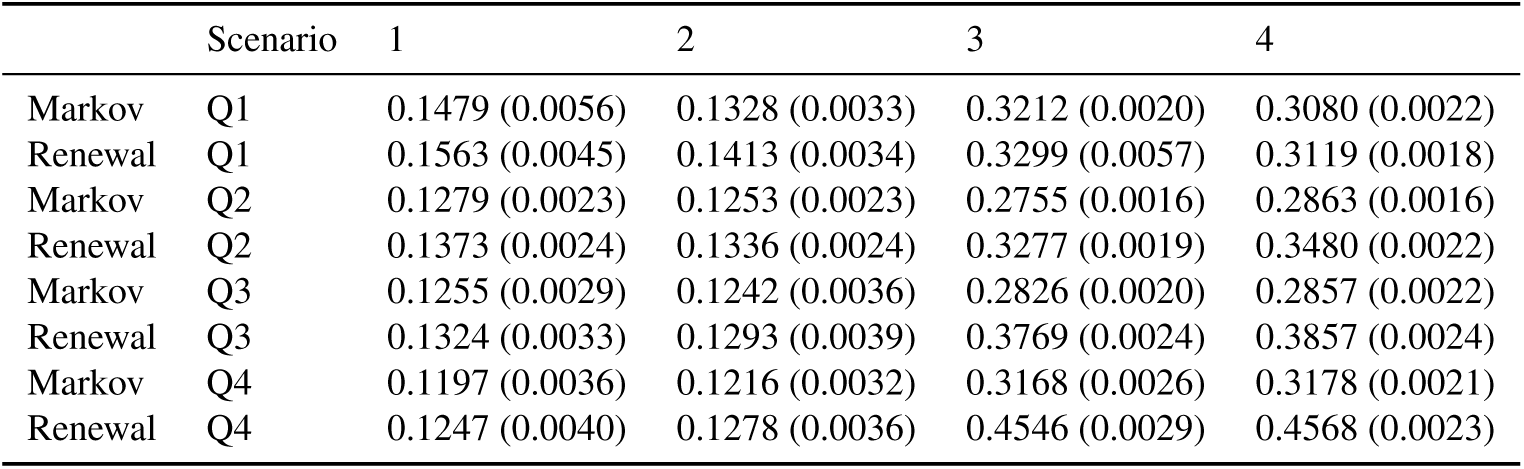
Mean relative error (Err_B_) over 25 runs under each scenario stratified by quartile. Standard error in parenthesis.

**Table S3.** Mean counts of loci in each quarter for under each scenario across 25 simulations. Standard error in parenthesis.

| Scenario | 1 | 2 | 3 | 4 |
| --- | --- | --- | --- | --- |
| Q1 | 25878.20 (951.37) | 28103.76 (1075.15) | 27807.96 (580.26) | 27659.64 (233.24) |
| Q2 | 12037.56 (646.81) | 10589.40 (702.38) | 10931.92 (331.94) | 11149.32 (143.83) |
| Q3 | 7917.24 (461.61) | 7231.24 (509.98) | 7200.76 (178.59) | 7119.12 (95.21) |
| Q4 | 4167.00 (360.71) | 4075.60 (272.09) | 4059.36 (124.89) | 4071.92 (90.72) |

**Table S4.**
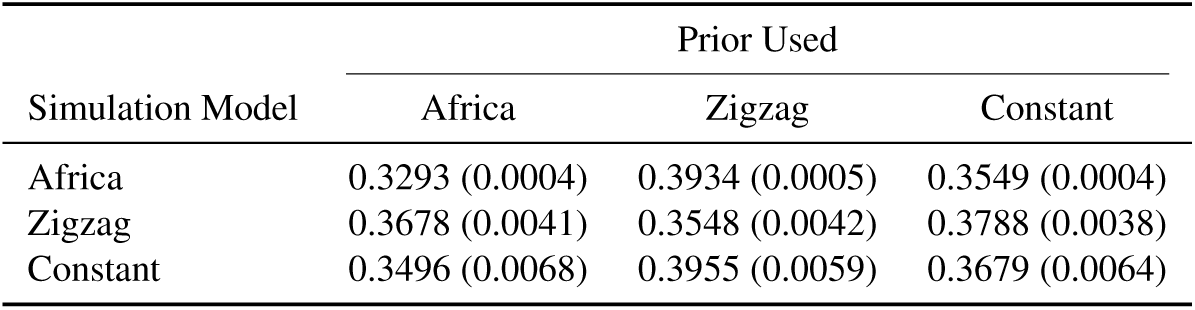
Mean relative error (Err_B_) over 25 runs under each scenario. Standard error in parenthesis.

| Simulation Model | Prior Used |  |  |
| --- | --- | --- | --- |
|  | Africa | Zigzag | Constant |
| Africa | 0.3293 (0.0004) | 0.3934 (0.0005) | 0.3549 (0.0004) |
| Zigzag | 0.3678 (0.0041) | 0.3548 (0.0042) | 0.3788 (0.0038) |
| Constant | 0.3496 (0.0068) | 0.3955 (0.0059) | 0.3679 (0.0064) |

**Table S5.**
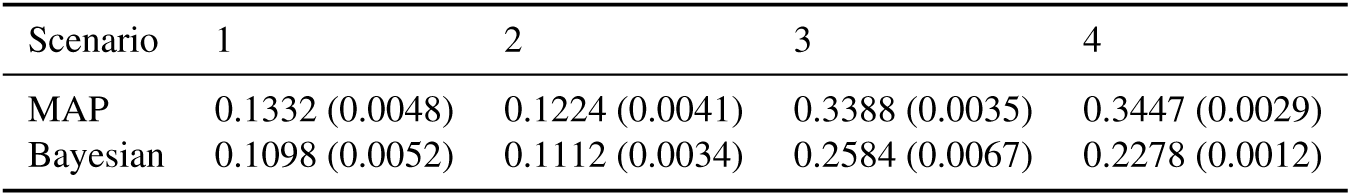
Mean relative error (Err_B_) over 25 runs under each scenario. Standard error in parenthesis.

| Scenario | 1 | 2 | 3 | 4 |
| --- | --- | --- | --- | --- |
| MAP | 0.1332 (0.0048) | 0.1224 (0.0041) | 0.3388 (0.0035) | 0.3447 (0.0029) |
| Bayesian | 0.1098 (0.0052) | 0.1112 (0.0034) | 0.2584 (0.0067) | 0.2278 (0.0012) |

**Fig S4.**
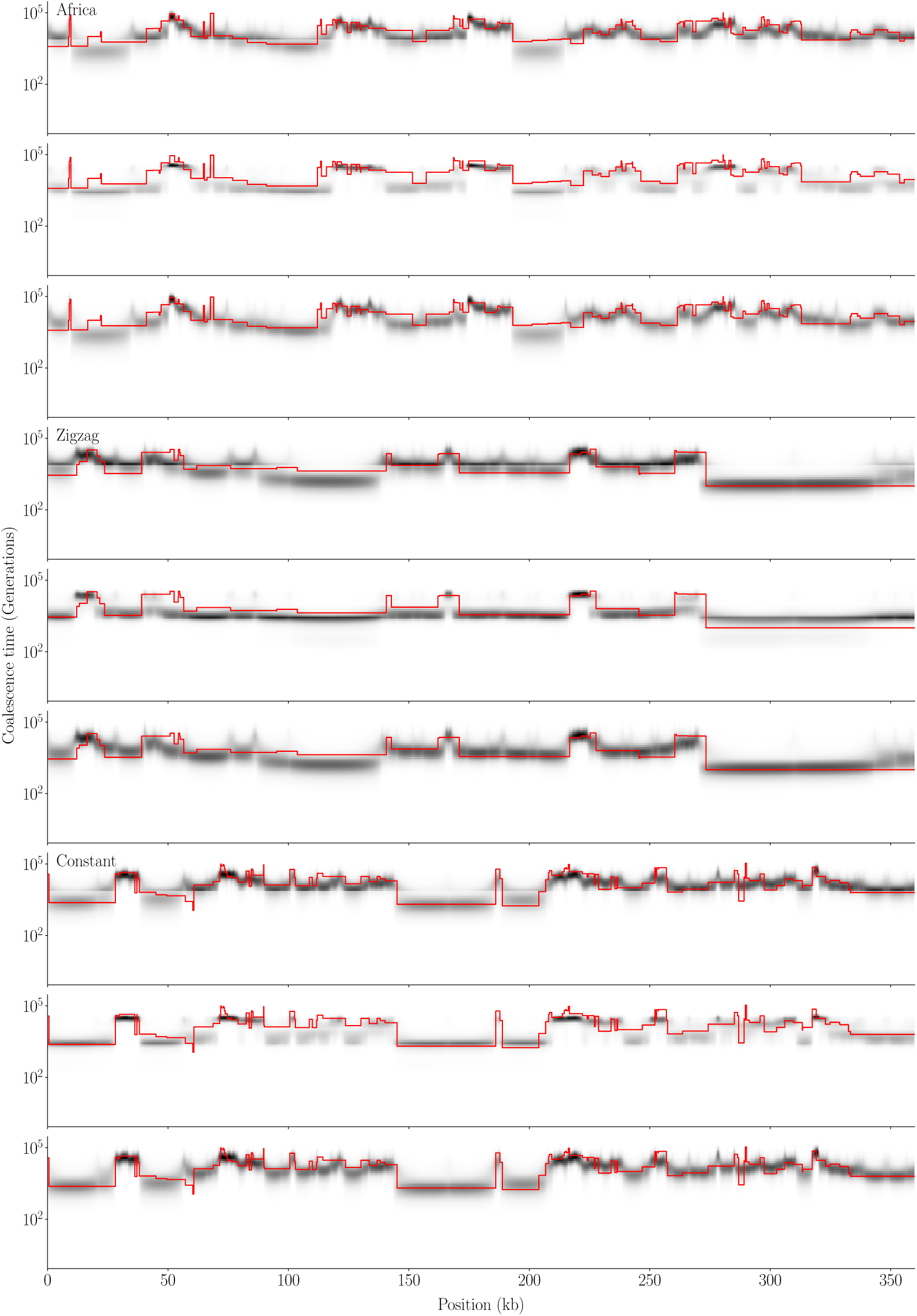
Comparison of posterior using different demographic priors. The figure is broken into groups of three panels where the data in the first three panels were generated by the Africa demography model, the second three by the zigzag model, and the last three by a constant population size model. Within each group of three, the first panel is the posterior using the Africa model as a demographic prior, the second using the zigzag model, and the last using a constant model. The red line in each panel is the true TMRCA.

**Fig S5.**
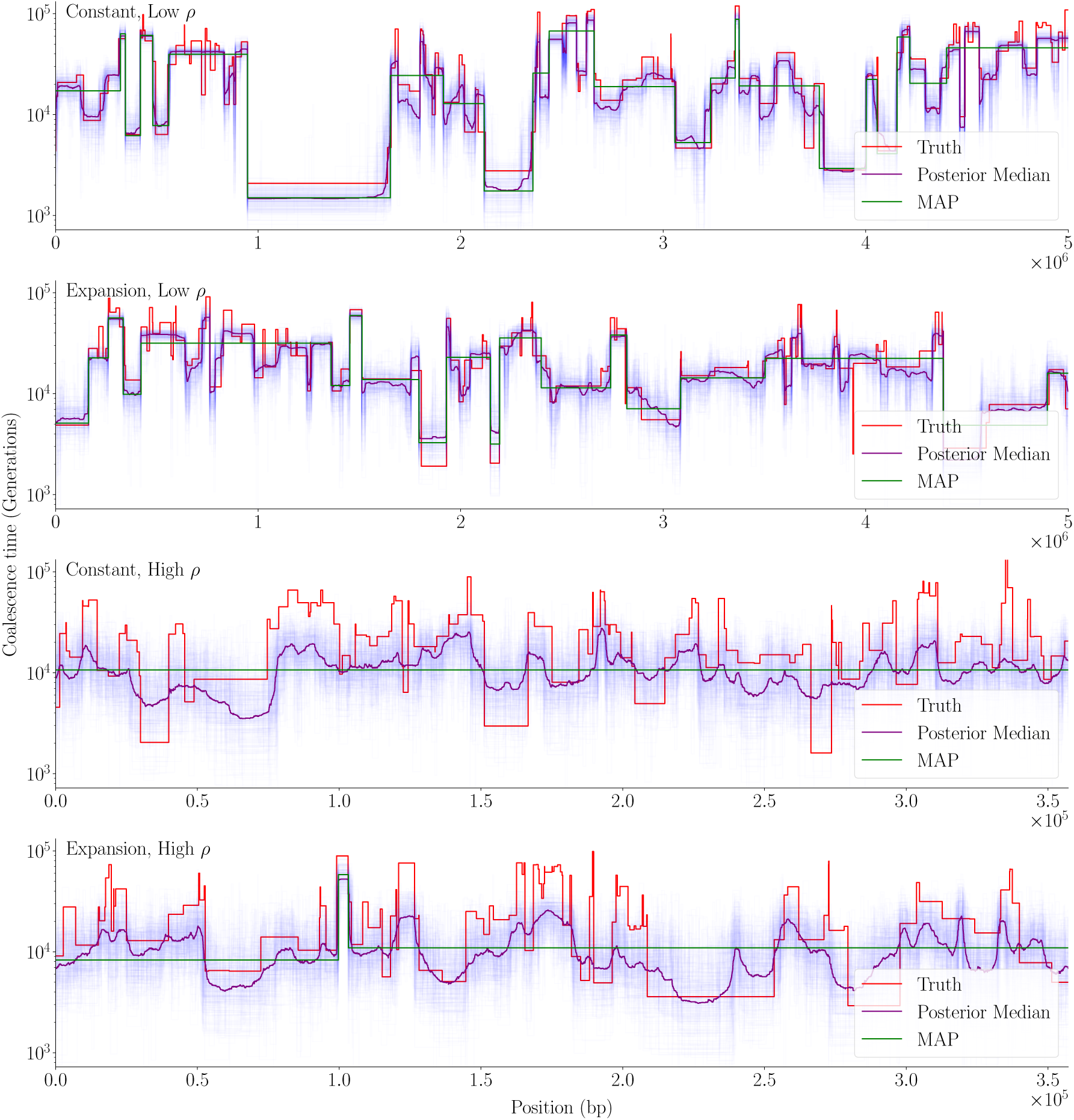
Comparison of Bayesian and frequentist method on simulated data. The light purple lines represent sample paths drawn from the posterior.

**Fig S6.**
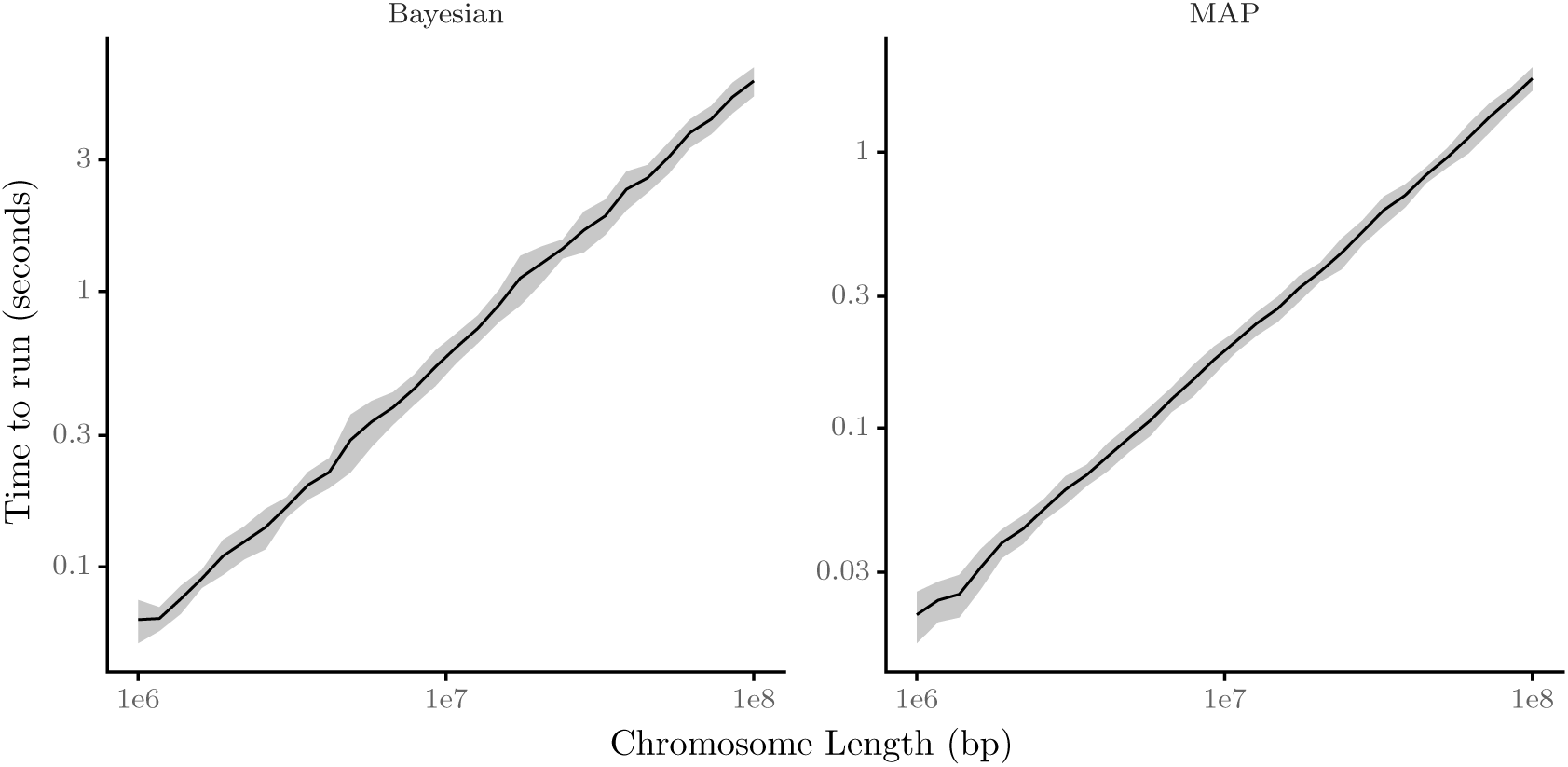
Mean running time of XSMC over various chromosome lengths on a log-log scale. The bands represent the standard error of the runs.

## Footnotes

1 This acronym is shared with a well-known sampling procedure in Bayesian statistics. In this paper, we only ever use SMC to refer to the sequentially Markov coalescent.

2 In the most granular analysis of sequence data, we can treat each nucleotide as an individual locus. However, if recombination is rare, then a large computational speedup can be obtained, with little effect on accuracy, by grouping nucleotides into windows of length e.g., 100, and assuming that recombination only occurs at the bound-aries between adjacent windows. For this reason, we describe our model generically in terms of non-recombining loci, rather than focusing specifically on sequence data.

